# Neural reactivation of past events tracks narrative understanding as it unfolds

**DOI:** 10.64898/2026.11.16.766768

**Authors:** Ka Ying Ivy Chan, Iman Mouloudi, Aya Ben-Yakov, Haim Dubossarsky, Avital Hahamy

## Abstract

Understanding of episodic experience is not achieved at a single moment: it is a constructive process in which each new event is interpreted in light of relevant past events and prior knowledge. Yet this process is rarely measured as it unfolds, leaving unclear how the brain integrates past events to construct an evolving understanding of the present. We hypothesized that neural reactivation of past events at narrative event boundaries reflects not merely what people experience, but how they interpret their experience. We developed a behavioural paradigm that tracks, as a narrative unfolds, which past events people consider relevant for understanding each new event. By manipulating prior knowledge of the narrative’s final plot twist, we examined the process by which the same experience yields different understanding. Across independent behavioural and fMRI cohorts, we found that prior knowledge similarly altered the variability and intersubject similarity of inferred relations between events and their neural reactivation at narrative event boundaries in the Default Mode Network. Moreover, stronger neural reactivation was associated with greater inferred relevance between events, and event pairs showing different behavioural relations across groups also showed corresponding differences in reactivation. These findings suggest that neural reactivation of past events at event boundaries reflects an unfolding inferential process that transforms shared experience into subjective understanding.

## Introduction

Understanding of real-world episodic experience is a constructive process, integrating many pieces of information over time. Yet, understanding is rarely studied as an *unfolding process* – it is typically reduced to its end-state: a consolidated memory, correct answer, or an “aha” moment. Through such approaches, seminal work has revealed that regions of the Default Mode Network (DMN, including the hippocampus) transform experience into knowledge^1–9^ and support moments of insight^10–16^.

Neuroimaging studies using movies or stories have further deepened our understanding of DMN involvement in insight^17–19^, and in the segmentation of experience into discrete events that are later integrated into knowledge^20–32^. This integration is often reflected in changes in fine-grained neural activity patterns, termed representations^33–36^. Yet, while naturalistic research has captured important neural dynamics that operate as experience unfolds, the lack of equally detailed behavioural measures leaves unclear how these dynamics reflect the process through which ongoing experience transforms into understanding.

One neural mechanism potentially relevant to the unfolding construction of understanding is the reactivation of past events. We have recently developed a method to detect such reactivation in naturalistic fMRI data^37^. Using this approach, we revealed that at transitions between discrete narrative events – commonly termed event boundaries^20–22^ – DMN regions reactivate representations of semantically-relevant past events, as if weaving them into an overarching narrative^37^ (and also see^38–41^). But does this neural reactivation simply reflect what people experience, or could it also reflect how they interpret their experience? Here we hypothesized that reactivation reflects the unfolding process of understanding, selectively linking past events to the present according to their subjective relevance.

To test this hypothesis, we directly manipulated the understanding of a naturalistic experience. We made use of an existing fMRI dataset^30^, in which participants watched the same movie either with or without prior knowledge of its final plot twist, such that understanding was either guided by a pre-existing interpretive framework or constructed in real time as the narrative unfolded. Building on this design, we developed a behavioural paradigm in which a new cohort of participants watched the same movie under the same manipulation. In this paradigm, after each episodic event, participants rated how relevant each previously-viewed event was for understanding the current one, allowing us to track the fine-grained inferential process through which understanding is constructed on the fly. We tested for group-differences in the unfolding process of understanding using two preregistered hypotheses, and found that prior knowledge produced changes in the variability and intersubject agreement of behavioural judgements, and that corresponding differences existed in the neural reactivation of past events. Moreover, we found that neural reactivation at event boundaries directly reflected the unfolding process of narrative understanding in both groups, and that event pairs showing group differences in inferred relations also showed corresponding differences in reactivation. This correspondence was evident across DMN regions and was stronger in the hippocampus when understanding was constructed in real time rather than guided by prior knowledge. Together, these findings demonstrate that the neural reactivation of past events at naturalistic event boundaries reflects not merely the structure of experience, but how people construct an evolving understanding of it.

## Results

We aimed to measure the behavioural process of narrative understanding, and test whether it is supported by the neural integration of events over time. To this end, we used an edited version of the film The Sixth Sense, whose plot culminates in a narrative twist^30^. Participants either watched the film naively, building their understanding of the narrative as it unfolded (Naïve condition), or were informed of the twist in advance, allowing them to interpret the narrative as it unfolded through a pre-existing interpretive framework (Spoiled condition). We measured the behavioural process of understanding by asking participants to provide real-time, scene-by-scene ratings, indicating how important each previously-viewed scene was to understanding each current scene, yielding ratings for all 666 scene pairs across the film (Fig. 1a, b). To measure the neural process of understanding, we made use of a freely available fMRI dataset in which independent groups of participants passively watched the same two versions of the film^30^. We measured the neural integration of experience over time through the reactivation of past scenes at movie event boundaries^37^ (Fig. 1c). We first characterized the behavioural and neural signatures of narrative integration separately, by examining differences in behavioural ratings and fMRI reactivation between the Naïve and Spoiled conditions. We then directly tested whether the behavioural process of understanding was reflected in fMRI reactivation of scenes at event boundaries.

**Figure 1.**
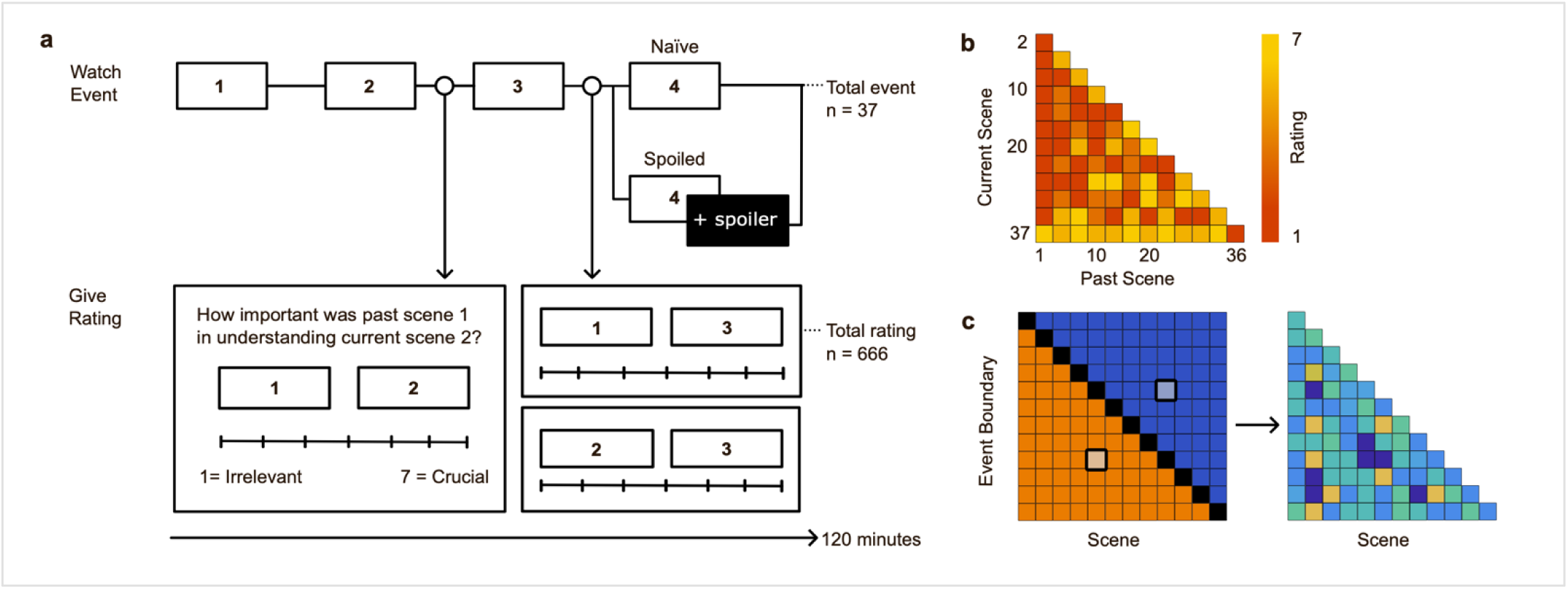
Experimental design and reactivation measurement schematics. (a) Participants watched a 60-minute film segmented into 37 scenes. At the end of each scene (open circles), they rated how important each preceding scene was for understanding the current scene on a scale from 1 (Irrelevant) to 7 (Crucial), resulting in 666 ratings per participant (experimental trials are illustrated in the bottom panel). Participants were assigned to one of two conditions: The Naïve group watched the original film with its twist ending. The Spoiled group received a written spoiler immediately after scene 4, revealing a fact that recontextualizes the interpretation of the rest of the film and explains its eventual twist ending. (b) Illustration of a behavioural rating matrix. Each cell represents the relational rating of a past scene (columns) for understanding a current scene (rows), with color indicating the rating magnitude (1 = dark red, 7 = yellow). (c) Schematic of neural reactivation matrices. fMRI pattern similarity was computed between the representation at each event boundary and the representation of each scene (left). In the resulting matrix, the lower triangular part contains similarities between each event boundary’s representation and the representations of preceding scenes (orange), whereas the upper triangular part contains similarities between each event boundary’s representation and the representations of subsequent scenes (blue). Each similarity value in the upper triangular part was subtracted from the corresponding value at its mirrored position in the lower triangular part (illustrated by the highlighted cells), resulting in a lower triangular matrix of the same size as the behavioural matrix (b), depicting the degree to which each scene representation is reactivated at each event boundary (right).

### The behavioural process of understanding

#### Inferred event relations capture the unfolding structure of narrative understanding

We first asked whether our novel relational-rating measure captured meaningful structure in participants’ evolving understanding of the narrative. For visualization, we constructed group-averaged rating matrices in which each cell represented the mean rating assigned to a given past–current scene pair across participants (Fig. 2a). Visual inspection verified that the resulting patterns reflected the narrative structure across multiple temporal scales. Short-range integration was demonstrated in the upper boundary of the lower-triangular matrices, which were assigned the highest mean ratings. This indicated that participants reliably judged the immediately preceding scene as important for understanding each current scene, consistent with the linear structure of the narrative. Intermediate-range integration was demonstrated in two clusters of higher ratings, reflecting strong inter-relations among scenes 13–20 and among scenes 28–33. These clusters correspond to two self-contained narrative subplots, each of which unfolded and resolved within its own scene range, suggesting that participants selectively inferred connections between events that belonged to the same underlying narrative thread. Long-range integration was also evident, as participants showed high ratings for the relation between the final, twist-revealing scene (37) and early scenes (1–4), whose meaning become clear only at the ending. Overall, these qualitative features indicate that the behavioural ratings captured the evolving inferential structure of the narrative, selectively linking events according to their narrative relevance. This relational structure was largely shared across conditions, as reflected in the highly similar group-averaged matrices (Spearman *r = 0.935*, *p < 0.001*).

**Figure 2.**
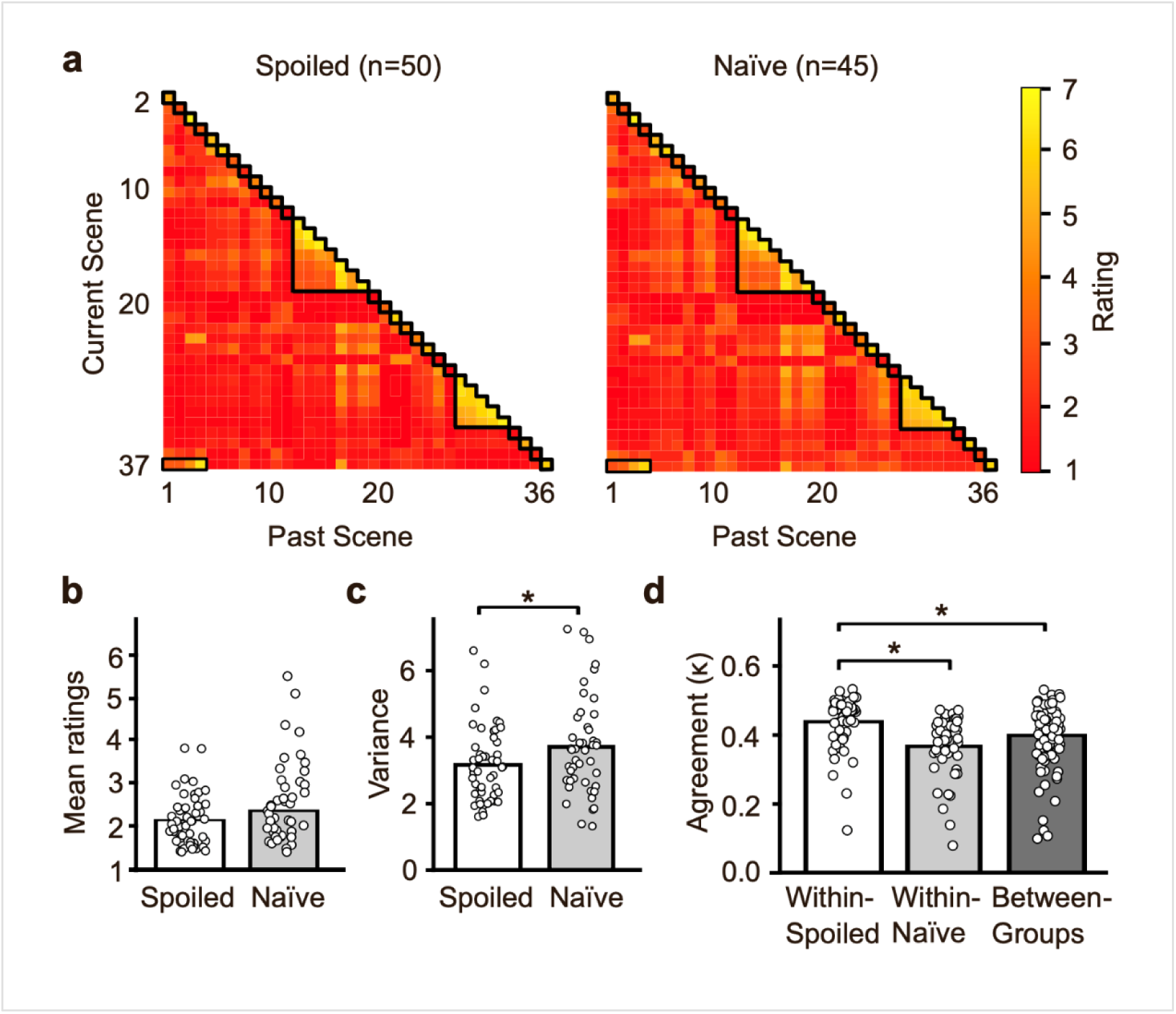
Behavioural measures of the unfolding process of understanding. (a) Group-averaged relational rating matrices for the Spoiled (*n = 50*) and Naïve (*n = 45*) conditions. Color indicates the mean rating (1 = dark red, 7 = yellow). Black contours highlight three features common to both matrices: (1) higher ratings along the upper boundary of the matrices, corresponding to the relation between the immediately preceding scene and the current scene; (2) two clusters of higher ratings among scenes 13–20 and among scenes 28–33, corresponding to self-contained narrative subplots; and (3) higher ratings linking the final scene (37) to the earliest scenes (1–4), reflecting integration of the twist with the film’s opening events. (b) Mean within-participant relational rating across all scene pairs. The Naïve group showed a non-significant trend toward higher ratings than the Spoiled group. (c) Within-participant variance of relational ratings across all scene pairs. The Naïve group showed significantly greater variance than the Spoiled group. (d) Intersubject agreement of ratings within and between groups, averaged across all participant-pairs for each participant. Within-Spoiled agreement significantly exceeded both between-group and within-Naïve agreement. Bars indicate group means; dots represent individual participants. *\*p < 0.05*.

#### Prior knowledge alters the distribution of relational ratings

We started by testing our first preregistered hypothesis that differences in prior knowledge of the twist would alter the distributions of individual participants’ relational ratings, as reflected in their means and variances. To this end, we first compared each participant’s mean rating across all scene pairs between the Naïve and Spoiled groups (Fig. 2b). The Naïve group showed slightly higher mean rating than the Spoiled group (Naïve: *M = 2.36*, *SD = 0.78*; Spoiled: *M = 2.11*, *SD = 0.53*), but this difference reached only trend-level significance (*p = 0.07*, two-tailed permutation test; see Supplementary Fig. 1b for analysis of median ratings).

We next compared the variance of ratings across all scene pairs for each participant (Fig. 2c). The Naïve group showed significantly greater rating variability compared to the Spoiled group (Spoiled: *M = 3.07*, *SD = 1.11*; Naïve: *M = 3.59*, *SD = 1.43*; *p = 0.045,* two-tailed permutation test; see Supplementary Fig. 1c for analysis of interquartile ranges), suggesting greater dispersion in judgements when the narrative’s interpretation was less constrained by prior knowledge.

#### Convergence of relational ratings across participants

Our next preregistered hypothesis was that the process of understanding, as reflected in the rating patterns, would be more consistent within conditions than between conditions, because prior knowledge was expected to shape the inferential relations participants formed between events. To test this, we quantified intersubject agreement by computing quadratic-weighted Cohen’s κ for every participant pair and compared these levels of agreement within and between groups (Fig. 2d). In line with our hypothesis, within-group agreement exceeded between-group agreement (within: *κ = 0.404*; between: *κ = 0.398*; *p = 0.022*, one-tailed permutation test). This effect was driven by the Spoiled group: Agreement among Spoiled participants was higher than between-group agreement (*p = 0.020*, one-tailed permutation test), and also higher than among Naïve participants (within-Spoiled: *κ = 0.438*; within-Naïve: *κ = 0.367*; *p = 0.047*, two-tailed permutation test). By contrast, agreement among Naïve participants did not exceed between-group agreement (*p = 0.965*; see Supplementary Fig. 1d for similar results obtained using Pearson correlations). Together, these findings suggest that prior knowledge promotes convergence in the behavioural process of narrative understanding. When viewers are given the narrative’s interpretive framework in advance, incoming events can be evaluated relative to that shared framework, leading to more convergent interpretations over time; without it, the same events may support multiple plausible interpretations, allowing individuals’ understanding to diverge as the narrative unfolds.

#### Subjective evaluation of the task

Post-experiment questionnaires confirmed that the two groups did not differ in their subjective evaluation of the task, including instruction clarity, engagement, task difficulty, or its perceived progression (all *p > 0.45*, two-tailed permutation tests; Supplementary Fig. 2). These findings suggest that differences in task experience are unlikely to contribute to the behavioural results reported above.

#### The neural process of understanding

We so far found that, compared to Spoiled participants, the Naïve participants showed significantly greater individual variability and a tendency toward overall higher relational ratings, consistent with greater uncertainty in narrative interpretation. In contrast, Spoiled participants showed greater agreement with one another, reflecting a shared interpretive framework imposed by prior knowledge. We next examined whether these behavioural differences were mirrored in the neural reactivation of past events at narrative event boundaries in an independent sample of participants, passively watching the same two versions of the movie^30^.

#### Reactivation of past scenes at event boundaries

We first replicated our previous findings of reactivation of past scene representations at event boundaries of naturalistic stimuli in DMN regions^37^. As demonstrated in Fig. 3, a whole-brain searchlight analysis revealed reactivation in both groups in the angular gyrus/lateral occipital cortex (ANG/LOC), precuneus/retrosplenial cortex (PCUN), and hippocampus (HPC; permutation-based cluster-mass inference with a cluster-forming threshold of *p < 0.001* and whole-brain family-wise error correction at *p < 0.05*), consistent with our previous findings^37^.

**Figure 3.**
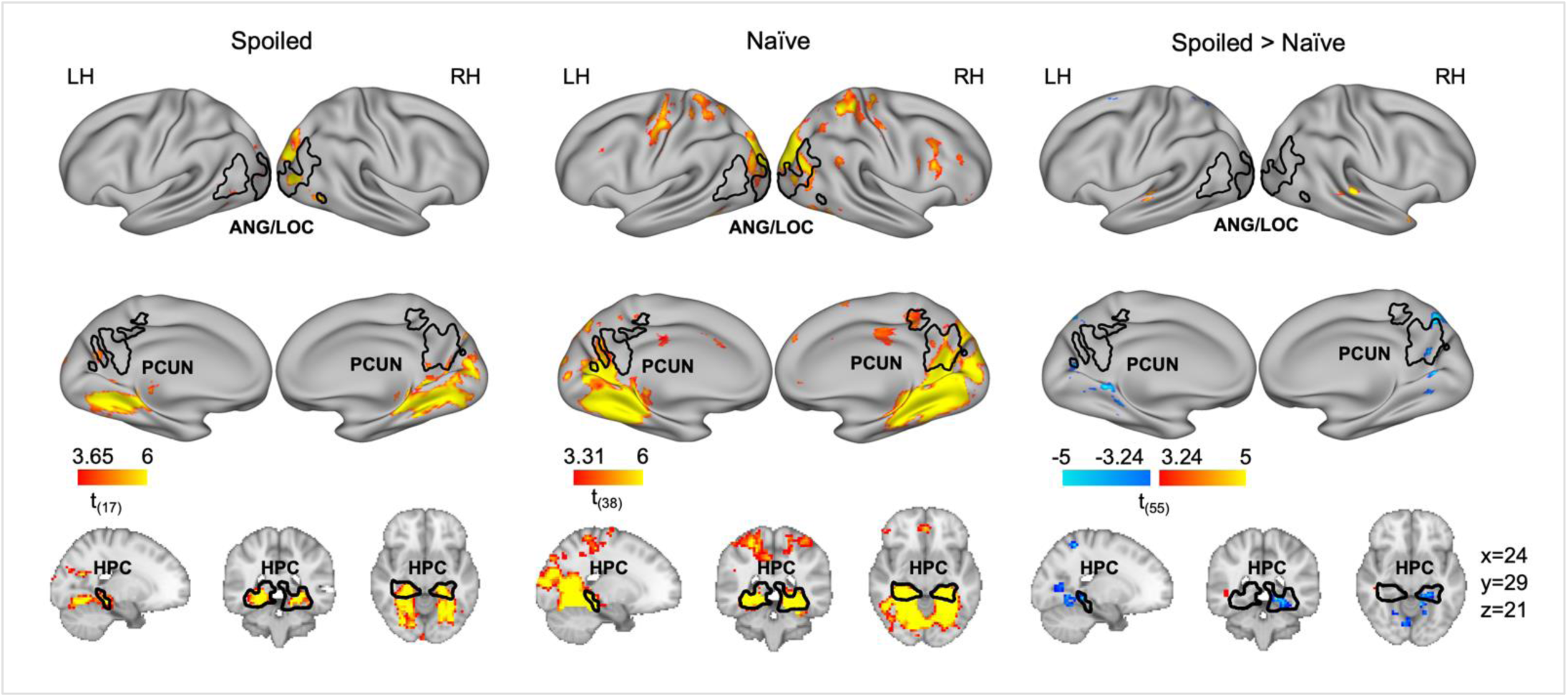
Whole-brain reactivation of past events at narrative event boundaries. Whole-brain searchlight reactivation maps for the Spoiled (left; *n = 18*) and Naïve (middle; *n = 39*) groups, and their between-groups contrast (right; warm colors indicate greater reactivation in Spoiled group, whereas cool colors indicate greater reactivation in Naïve group). Maps are presented on both flat cortical surfaces and on three-dimensional slices (dataset-specific MNI coordinates are depicted). Both groups showed reactivation of past events (positive t values) in the bilateral precuneus/retrosplenial cortex (PCUN), Angular Gyrus/Lateral Occipital Cortex (ANG/LOC) and hippocampus (HPC). Compared with the Spoiled group, the Naïve group showed greater reactivation in the PCUN and HPC. LH, left hemisphere; RH, right hemisphere. Black contours outline independently defined ROIs from Hahamy et al.^37^. Within-group maps were obtained using one-tailed one-sample t-tests; the between-group contrast used a two-tailed two-samples t-test. All maps were corrected for multiple comparisons using permutation-based cluster-mass inference (cluster-forming threshold p < 0.001; whole-brain FWE-corrected *p < 0.05*). See Supplementary Table 1 and 2 for complete cluster statistics.

An independent ROI analysis confirmed these observations. Both the Spoiled and Naïve groups showed significant reactivation of past events at event boundaries in a composite DMN ROI (Spoiled: *p < 0.001*; Naïve: *p = 0.007*, one-tailed permutation tests). These effects also held in each separate ROI (all p ≤ 0.006, one-tailed permutation tests, Holm-Bonferroni corrected; Fig. 4a).

**Figure 4.**
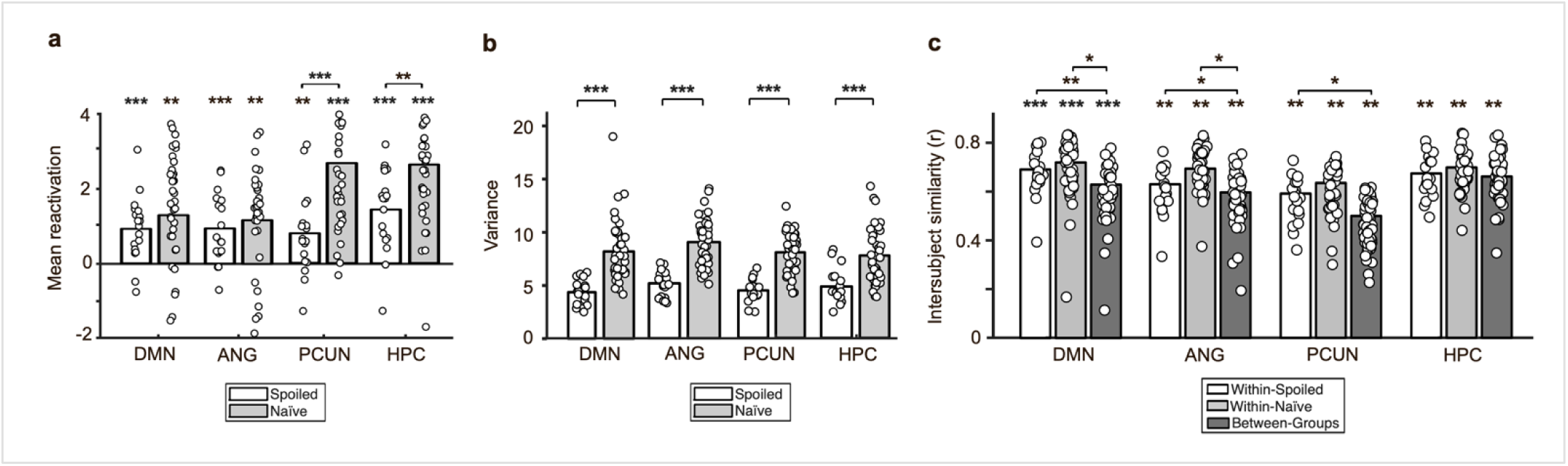
Neural measures of the understanding process. (a) Mean reactivation index for participants in the Spoiled (white bars; *n = 18*) and Naïve (gray bars; *n = 39*) groups across the composite Default Mode Network (DMN) ROI, angular gyrus/lateral occipital cortex (ANG), precuneus/retrosplenial cortex (PCUN), and hippocampus (HPC). (b) Variance of the granular reactivation patterns for each participant, shown for the same ROIs and groups. (c) Intersubject similarity of the granular reactivation pattern for the Spoiled (white), Naïve (light gray), and between-group (dark gray) comparisons. Bars indicate group means; dots represent individual participants. For the exploratory ROI analyses (ANG, PCUN, and HPC), p-values were Holm-Bonferroni corrected. Asterisks above bars indicate that the corresponding group mean differs significantly from zero; bracket asterisks indicate significant between-group differences (permutation tests). *p < 0.05, **p < 0.01, ***p < 0.001.

Although reactivation was also found in additional regions (see Supplementary Table 1 for full cluster statistics), we focus subsequent analyses on these predefined DMN regions, as they had been previously identified as most replicable across independent datasets^37^.

#### Prior knowledge alters the distribution of reactivated scenes

Visual inspection of a whole-brain between-group comparison revealed that the Naïve group showed greater reactivation than the Spoiled group in the precuneus and hippocampus (permutation-based cluster-mass inference with a cluster-forming threshold of *p < 0.001* and whole-brain family-wise error correction at *p < 0.05*; Fig. 3, right panel; see Supplementary Table 2 for full cluster statistics).

In line with our behavioural results, reactivation in the composite DMN ROI showed a tendency toward higher reactivation in the Naïve compared to the Spoiled group, which was not significant (*p = 0.763*) (Fig. 4a). Breaking this effect into specific ROIs confirmed significantly greater reactivation for the Naïve group in the precuneus (*p < 0.001*) and hippocampus (*p = 0.007*), but not in angular gyrus (*p = 0.807*, all two-tailed permutation tests, Holm-Bonferroni corrected).

We further compared participants’ variance in granular reactivation values between groups (Fig. 4b). Consistent with the behavioural findings, the Naïve group showed significantly greater variance than the Spoiled group in the DMN ROI (*p < 0.001*, two-tailed permutation test). This effect was also observed across all ROIs (all *p < 0.001*, two-tailed permutation tests, Holm-Bonferroni corrected; see Supplementary Table 3 for means, SEMs, and full statistics), indicating more dispersed reactivations when the narrative’s interpretation is more uncertain.

#### Convergence of reactivated scenes across participants

To test whether shared prior knowledge led to more consistent neural reactivation patterns within compared to between conditions, we measured the intersubject similarity of the granular reactivation patterns across all participants (Fig. 4c). In line with our behavioural results, within-group reactivation similarity exceeded between-group similarity in the DMN ROI (within: *M = 0.705*; between: *M = 0.629*; *p = 0.007*, one-tailed permutation test). Unlike the behavioural results, both within-Spoiled and within-Naïve similarity individually exceeded between-group similarity (within-Spoiled: *M = 0.690*; within-Naïve: *M = 0.719*; between: *M = 0.629*; both p ≤ 0.024, one tailed permutation tests), and no group difference was found in the within-group similarities (within-Spoiled vs. within-Naïve: *p = 0.509*, two-tailed permutation test).

Further decomposing the DMN into separate ROIs revealed that in the angular gyrus, within group exceeded between group similarities in both groups (within-Spoiled: *M = 0.630*; within-Naïve: *M = 0.694*; between: *M = 0.596*; both p ≤ 0.033, one-tailed permutation test, Holm-Bonferroni corrected). In the precuneus, within-group similarity exceeded between-group similarity for the Spoiled (*p = 0.024*, one-tailed permutation test, Holm-Bonferroni corrected) but not the Naïve group (*p = 0.154*, one-tailed permutation test, Holm-Bonferroni corrected). No significant effects were detected in the hippocampus (within-Spoiled vs. between: *p = 0.605*; within-Naïve vs. between: *p = 0.985*; one-tailed permutation test, Holm-Bonferroni corrected). The two within-group similarities did not differ in any ROI (all p ≥ 0.666, two-tailed permutation test, Holm-Bonferroni corrected).

These neural results broadly mirror our behavioural findings. While we found significant DMN reactivation of past scenes at event boundaries in both groups, the Naïve participants showed a non-significant tendency for stronger reactivation compared to the Spoiled participants, with significant group differences in the precuneus and hippocampus. The Naïve group also showed greater variability in reactivation patterns across the DMN, consistent with the greater interpretive uncertainty associated with the Naïve condition. Finally, within-group reactivation patterns were generally more similar than between-group patterns, indicating condition-specific neural signatures of narrative understanding.

#### Neural reactivation tracks the behavioural process of understanding

Having found parallel behavioural and neural signatures of understanding, we next asked whether the two were directly related. To test this, we correlated neural reactivation with behavioural relational ratings within each experimental condition (Fig. 5a; see also Supplementary Fig. 1e and 3). Both groups showed significantly positive brain-behaviour correlations in the DMN ROI (both *p < 0.001*, one-tailed permutation tests). This relationship was also significant in each individual ROI (all *p < 0.001*; Holm-Bonferroni corrected, one-tailed permutation tests), indicating that scene pairs judged as more important for understanding were associated with stronger neural reactivation, and vice versa.

**Figure 5.**
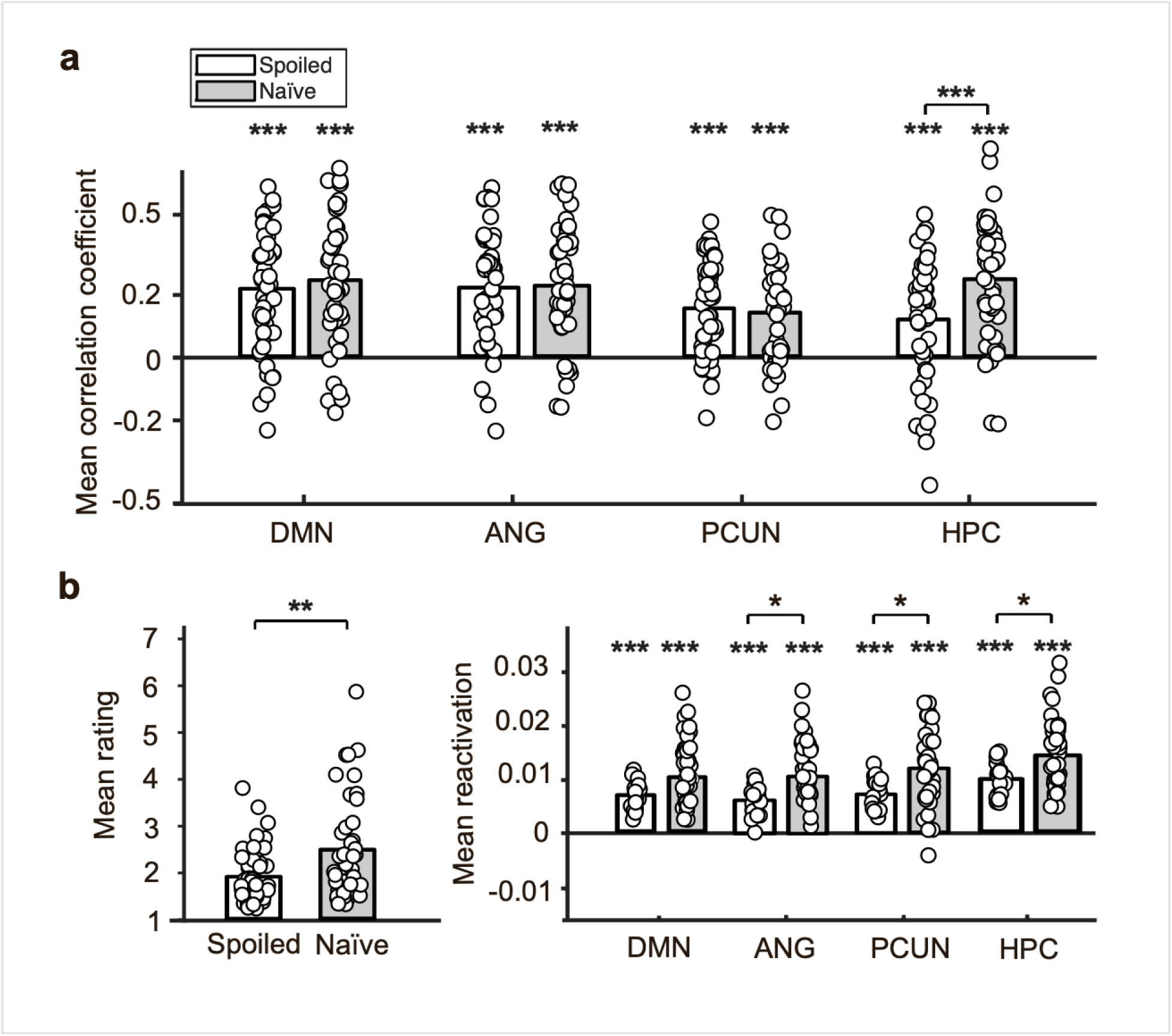
Brain–behaviour correspondence. (a) Correlation coefficients relating behavioural rating level (1–7) to granular reactivation patterns, computed for each same-condition behavioural-fMRI participant pair. Correlations are shown separately for the Spoiled (white bars; behavioural *n = 50*) and Naïve (gray bars; behavioural *n = 45*) groups. (b) Exploratory analysis testing whether the top 25% of scene pairs showing the largest behavioural differences between groups also exhibited corresponding differences in neural reactivation. Left: group differences in the mean behavioural rating across the selected scene pairs. Because the scene pairs were selected on the basis of this behavioural difference, the bracket asterisks on this panel reflect the magnitude of the selection criterion itself rather than an independent statistical test, and are shown for reference only. Right: Mean reactivation across the selected scene pairs for each ROI, showing greater neural reactivation for the selected scene pairs in the Naïve group than in the Spoiled group in angular gyrus, precuneus, and hippocampus, with a non-significant trend in DMN (two-tailed permutation tests). Bars indicate group means; dots represent individual participants. Asterisks above the bars indicate that the corresponding group mean differs significantly from zero; bracket asterisks indicate significant between-group differences (permutation tests). For the exploratory ROI analyses (ANG, PCUN, and HPC), p-values were Holm-Bonferroni-corrected. *\*p < 0.05, **p < 0.01, ***p < 0.001*. See Supplementary Fig. 1e for complementary correlation analyses (panel a), and Supplementary Table 6 for analyses using alternative selection thresholds (panel b).

Between-group comparisons of brain-behaviour coupling did not differ in the DMN, angular gyrus, or precuneus (all p ≥ 0.434). In contrast, coupling was significantly stronger in the hippocampus for the Naïve than the Spoiled group (Spoiled: *M = 0.128*, *SD = 0.208*; Naïve: *M = 0.268*, *SD = 0.209*, *p < 0.001*, Holm-Bonferroni corrected, two-tailed permutation test).

To assess whether this hippocampal group difference reflected differences in the overall mean or variance of behavioural ratings or neural reactivation reported above, we repeated the analysis while controlling for participants’ mean and variance in both the neural and behavioural measures. The between-group effect remained significant when controlling for neural covariates (*β = +0.107*, *p = 0.002*) and behavioural covariates (*β = +0.119*, *p = 0.005*, two-tailed permutation tests). These results were robust to whether brain-behaviour correlations were averaged across fMRI participants to yield one value per behavioural participant, or across behavioural participants to yield one value per fMRI participant; both approaches produced the same pattern of between-group effects (see Supplementary Table 4). Together, these findings indicate that the hippocampal group difference reflects brain-behaviour coupling itself, rather than differences in the overall distributions of the neural or behavioural measures.

Because the two groups shared much of the narrative’s relational structure (*r = 0.935* between group-averaged behavioural ratings), whole-matrix brain-behaviour correlations could be dominated by these shared relations, potentially obscuring differences associated with prior knowledge. Indeed, correlating behavioural ratings with neural reactivation patterns from participants in the other experimental group replicated the within-group brain–behaviour coupling, with no significant difference in within- and cross-group correlations in any ROI (see Supplementary Table 5). The fact that brain-behaviour coupling was independently significant in both groups indicates that this relationship is both replicable and invariant across conditions. However, this whole-matrix analysis was insensitive to the relatively small subset of scene pairs on which the groups differed behaviourally. We therefore asked whether those specific scene pairs also exhibited differences in neural reactivation. To this end, we selected the top 25% of scene pairs with the largest absolute between-group differences in behavioural ratings (see Supplementary Table 6 for analyses using alternative selection thresholds). As expected, the Naïve group assigned higher ratings than the Spoiled group within this subset (Spoiled: *M = 1.89*, *SD = 0.56*; Naïve: *M = 2.47*, *SD = 1.07*; Fig. 5b, left panel). We then examined neural reactivation for these same scene pairs in the independent fMRI cohort. Neural reactivation for these scene pairs was significantly stronger in the Naïve group than in the Spoiled group in the angular gyrus, precuneus, and hippocampus (all *p = 0.028*, Holm-Bonferroni-corrected), with a non-significant trend in the composite DMN ROI (*p = 0.075*, Fig. 5b, right panel).

Because the Naïve group also showed higher reactivation overall (Fig. 4a), we tested whether this effect was specific to the selected pairs by comparing, for each participant, reactivation for these pairs with reactivation for the remaining, non-selected pairs (see Methods). In both groups, and across all regions, reactivation was significantly greater for selected than for non-selected pairs (effect sizes ranged from *d = 1.48–2.94* across ROIs and groups, p < 0.001; Holm-Bonferroni-corrected). Critically, this selected-versus-non-selected reactivation difference was significantly greater in the Naïve than the Spoiled group in the DMN (*p = 0.018*), angular gyrus (*p = 0.004*) and hippocampus (*p = 0.035*), and showed a trend in the precuneus (*p = 0.053*; Holm-Bonferroni-corrected, see Supplementary Table 6 for analyses using alternative selection thresholds). These findings suggest that the localized behavioural group differences induced by uncertainty in narrative understanding are reflected in corresponding differences in neural reactivation that are not explained by the groups’ overall difference in reactivation.

## Discussion

What are the neural mechanisms of episodic understanding? Here, we studied understanding not as an end point, but as an unfolding inferential process. We replicated our previous finding of past-event reactivation at narrative event boundaries in the DMN, and further showed that this neural process mirrors the unfolding behavioural process of understanding. First, behavioural ratings and neural reactivation in the DMN showed parallel effects of prior knowledge: Naïve viewers, who lacked prior knowledge of the narrative’s resolution, showed greater variability in both measures together with higher mean values that reached significance only for neural reactivation in the precuneus and hippocampus. Second, intersubject convergence showed a similar overall pattern across behaviour and brain, with within-group consistency exceeding between-group consistency. Third, behavioural ratings and neural reactivation were significantly coupled in both groups, with stronger hippocampal coupling in Naïve than Spoiled viewers. Finally, scene pairs showing between-group differences in relational judgements also showed corresponding differences in neural reactivation. Importantly, these correspondences were observed in both experimental groups, despite the independent samples performing different tasks (passive movie viewing in the scanner versus deliberate inferential judgements in the behavioural experiment), suggesting a robust and reproducible link between neural reactivation and the unfolding construction of narrative understanding.

These findings extend previous demonstrations that prior knowledge and interpretive context influence the neural responses to naturalistic narratives^29, 42^ and the neural representations of narrative events^30, 31, 43, 44^. Our findings suggest that prior knowledge also guides inferences about which past events are relevant for understanding the present, thereby structuring the evolving relational organisation of experience. We demonstrate that prior knowledge changes the distributional properties of behavioural relations and neural reactivation across events. While mean values were generally higher in the Naïve group, the difference was non-significant behaviourally and regionally selective neurally, with significantly greater reactivation in the precuneus and hippocampus. By contrast, variance differed more consistently: Naïve participants showed a broader spread of values across scene pairs in both their behavioural ratings and neural reactivation. A plausible interpretation is that, without a fixed interpretive framework, the relevance of individual past events must be continually re-evaluated as new information arrives. Constructing understanding under uncertainty may therefore produce larger differences across event pairs: some past events become highly relevant when a new scene changes their interpretation, while others may become less relevant or remain unchanged. Prior knowledge, by contrast, provides a stable framework that constrains these evolving relations and may reduce the extent to which individual events are differentially weighted.

Despite the broad parallels between the behavioural and neural measures, one divergence emerged. Neural reactivation patterns were more similar within than between conditions for both Spoiled and Naïve participants, with comparable within-group similarities across the two conditions. Behaviourally, however, condition-specific agreement was evident only in the Spoiled group: Spoiled participants agreed more with one another than with Naïve participants, whereas Naïve participants showed lower levels of agreement that did not exceed the between-group agreement. One possible explanation is that Naïve participants in the fMRI dataset were screened to ensure that they had not inferred the twist before its formal reveal^30^, whereas participants in the behavioural study were not. If some behavioural Naïve participants guessed the twist early, their ratings may have shifted toward the Spoiled interpretation for parts of the narrative, reducing within-group agreement while increasing agreement across groups.

We also found a regional difference in how prior knowledge affected brain-behaviour coupling: hippocampal reactivation tracked inferential ratings more strongly in Naïve than Spoiled viewers, whereas coupling in cortical DMN regions was comparable across conditions. This effect was not explained by group differences in the mean or variance of either measure, indicating that prior knowledge altered the more fine-grained correspondence between hippocampal reactivation and behavioural understanding. These findings are consistent with proposals that the hippocampus supports the flexible formation and updating of relations among raw elements of experience^2, 3, 5, 6, 41^. When understanding is constructed under uncertainty (Naïve condition), the fine-grained process of relating each new event to specific past events, as measured in our task, may therefore more closely resemble hippocampal processing.

We have previously demonstrated the reactivation of past events at narrative event boundaries in DMN regions across two independent naturalistic datasets^37^ (and see also^19, 26, 38–41^). Here, we replicated this effect in two additional groups viewing a different naturalistic stimulus, with both Spoiled and Naïve participants showing reactivation in the same DMN regions. Importantly, while our previous work established that reactivation is selective for semantically relevant past events, the current study links this neural process to a direct behavioural measure of evolving understanding. Across all examined DMN regions, reactivation was stronger for past events that participants judged as more important for their subjective understanding of the present, and this relationship replicated across both experimental conditions. Furthermore, groups experiencing the same narrative through different interpretive frameworks showed corresponding differences in inferred event relations and neural reactivation for those event pairs. These findings suggest that reactivation does not merely reflect the structure of the narrative, but also the process by which its subjective interpretation is constructed.

More broadly, this study illustrates how understanding can be studied not only as an outcome, but as a process that unfolds over time. By combining a naturalistic narrative with continuous relational ratings, we were able to track how past events are progressively integrated into an evolving interpretation of the present, and to show that this behavioural process is reflected in neural reactivation. This process-level approach reveals structure that cannot be directly captured by conventional end-point measures or timepoint-focused approaches, and provides a framework for understanding how the brain extracts meaning from naturally unfolding experience.

## Method

This study combined a behavioural experiment with analysis of a freely available fMRI dataset^30^, both using the same stimulus: an edited version of The Sixth Sense (M. Night Shyamalan, 1999)^45^, watched under one of two conditions: Spoiled, in which participants were informed of the film’s twist ending at scene 4, or Naïve, in which they were not. The online behavioural experiment captured participants’ ongoing inferential process as they watched the movie, while the fMRI dataset provided neural reactivation measures from participants passively viewing the same stimuli. Together, these data allowed us to test whether the behavioural process of narrative understanding is reflected in neural reactivation of past events at narrative event boundaries.

### Experimental Stimulus

The experimental stimuli used here were developed by Zadbood et al.^30^, and comprised of edited versions of the movie The Sixth Sense^45^. Participants were assigned to one of two conditions, Spoiled or Naïve, which diverged in how the film’s twist was revealed. In the Spoiled condition, participants watched a ∼60-minute version of the film in which, at the end of scene 4, a textual spoiler revealed the film’s twist. This text was displayed against a black background with accompanying background music. In the Naïve condition, participants watched one of two versions preserving the film’s natural progression without foreknowledge of the twist: a ∼60-minute version, matched in length and pacing to the Spoiled version, but with the spoiler text replaced by a blank screen (with the original audio intact, preserving temporal alignment between the two conditions) and the twist was revealed naturally in the final scene; or a 55-minute version in which the final scene was removed entirely, such that the twist was never revealed. Apart from the above differences, all versions were identical in content, editing, and pacing across conditions. The experimental stimulus was segmented into events based on the agreement of two independent raters from the research team. Event boundaries mostly corresponded to scene cuts, marked by brief black-screen transitions between consecutive scenes. A small number of boundaries (*n=3*) did not coincide with scene cuts, but instead reflected the perceived ending of a narrative unit (e.g., the clear conclusion of a plot thread). Overall, 37 narrative events were identified, resulting in 666 unique event pairs.

### Behavioural Study

The behavioural experiment was preregistered prior to data collection^46^. We hypothesized that: 1) differences in prior knowledge of the twist would alter the distributions of individual participants’ relational ratings, as reflected in their means and variances; and 2) that participants within the same condition (Spoiled or Naïve) would show greater similarity in their relational ratings than participants across conditions, reflecting a shared narrative interpretation shaped by prior knowledge of the twist.

### Participants

We recruited 110 participants through Testable Minds, an online platform for behavioural research^47^. Eligibility criteria included native English speakers aged 18–50 who had not previously watched the experimental stimulus. Participants were excluded if they failed more than three of ten catch trials (*n = 8*), or assigned low ratings (<4 of 7) to the relation between the spoiler and twist-ending scene pairs in the Spoiled condition (*n = 7*), indicating that they had not integrated the spoiler with the twist ending, and hence did not reach the expected end-point understanding of the movie. A final sample of 95 participants was included in the analysis (46 females, 1 other; mean *age = 31.46 years*, *SD = 7.26*, *range = 22–50*).

Participants were randomly assigned to either the Spoiled (*n = 50*) or Naïve (*n = 45*) condition. The two groups were well-matched, with no significant demographic differences in age (*U = 1305, p = 0.180*), gender (*χ²(2, N = 95) = 1.52, p = 0.467*), or education level (*χ²(3, N = 95) = 2.42, p = 0.491*). All participants provided written informed consent prior to the experiment and received information about the conditions of the experiment and their rights, and were reimbursed for their time. The experiment protocol and the consent forms were approved by the Institutional Review Board of the Institute of Cognitive Neuroscience, UCL.

### Experimental Design

Participants watched the experimental stimuli, which was segmented into 37 narrative scenes. Each trial began at the end of a scene, where two representative screenshots were presented, one from the just-viewed (current) scene and one from a past scene. Participants were asked to rate how important the past scene was for understanding the current scene on a 7-point scale (1 = Irrelevant, 7 = Crucial). The comparison structure was incremental: ratings began at scene 2, which was rated against scene 1; each subsequent scene was then rated against every scene viewed so far (e.g., scene 3 was compared with scenes 1 and 2; scene 4 with scenes 1–3, and so on), with a cumulative timeline of previously viewed screenshots displayed beneath each pair to support in-context memory. This incremental structure yielded 666 pairwise relational ratings per participant.

Before the main experiment, all participants completed a two-phase tutorial to familiarize them with the task. In the demonstrative phase, participants passively viewed representative images of three scene pairs from the movie Little Red Riding Hood (1995)^48^, along with suggested ratings spanning the range from clearly unrelated to clearly essential relations, to calibrate their use of the full rating scale before the main task. For example, one pair showed Little Red Riding Hood picking flowers, followed by a later scene in which she asks, “Grandma, why are your teeth so big?”; participants were shown a suggested rating of 1, with the rationale that the flower-picking scene had no bearing on understanding the later exchange. In this phase, participants were not required to respond themselves, but were given the suggested rating and its accompanying explanation for each scene pair. In the interactive phase, participants watched and actively rated four scenes from the movie Snow White and the Seven Dwarfs (1937)^49^, using the same rating scale and incremental comparison structure as the main task (yielding six pairwise ratings). Each response was followed by a suggested rating and its explanation. Mean tutorial ratings reflected the expected narrative structure, with no group differences for any training pair (all *p > .60*, two-tailed Mann-Whitney U tests, Holm-Bonferroni-corrected).

To verify the participants’ attentiveness, ten catch trials were embedded at randomly determined, fixed positions throughout the experiment. In these trials, the earlier scene featured frames from other films starring the lead actor of the experimental stimulus, making them visually similar to the experimental stimulus yet irrelevant to the narrative plot. Participants were expected to assign a rating of “1” (irrelevant) to these trials. Mean catch trial ratings were *1.05 (SD = 0.13)* in the Spoiled condition and *1.05 (SD = 0.14)* in the Naïve condition, with no significant difference between groups (*p = .837*, two-tailed permutation test).

A post-task questionnaire assessed four items on a 7-point scale: clarity of instructions, movie engagement, overall task difficulty, and perceived increase in task difficulty.

### fMRI data

We made use of a freely available fMRI dataset^30^, consisting of 57 participants who passively watched the same edited versions of The Sixth Sense as in the behavioural experiment, originally divided into three groups: Spoiled (*n = 18*), who received a textual spoiler early in the movie; Twist (*n = 19*), who learned the plot twist only at the final scene; and No-Twist (*n = 20*), who watched a version with the twist edited out entirely. Zadbood et al.^30^ found that intersubject pattern similarity in the DMN during movie-viewing was significantly greater between the Twist and No-Twist groups than between either of these groups and the Spoiled group. This finding is expected from a narrative standpoint: until the final reveal, the Twist group (who learn the twist only at the very end) and the No-Twist group (who never encounter it) share the same interpretation of the unfolding narrative, unlike the Spoiled group, who interprets the narrative through knowledge of the twist from the outset. Therefore, for all our neural analyses, we pooled the Twist and No-Twist groups into a single Naïve group (*n = 39*).

### fMRI Acquisition

fMRI was performed on a 3T full-body Siemens Prisma scanner with a 64-channel head coil. Functional images were acquired using an interleaved multiband echo planar imaging (EPI) pulse sequence (TR: 1500 ms, TE: 33 ms, flip angle: 80°, multiband factor: SMS = 4), with each volume providing whole-brain coverage at a 2 mm slice thickness and FOV 192 mm². Anatomical images were acquired using a T1-weighted magnetization-prepared rapid-acquisition gradient echo (MPRAGE) pulse sequence with 1 mm³ resolution. Additionally, field maps were collected for B0 correction at the end of the functional runs.

### fMRI Preprocessing

Functional and anatomical MRI data were preprocessed using fMRIPrep 24.1.1^50^. Anatomical images were corrected, skull-stripped, and nonlinearly registered to the MNI152 template using fMRIPrep’s standard pipeline. For each functional run, slice timing correction and head motion correction were applied. Functional volumes were co-registered to the T1-weighted reference and normalized to MNI152 space. Framewise displacement (FD) and standardized DVARS were computed per run; volumes exceeding FD > 0.5 mm or DVARS > 1.5 were flagged as motion outliers. Nuisance signals were regressed from the functional time series, including six motion parameters, four anatomical CompCor components derived from a CSF mask, high-pass filtering regressors (300s cut-off), and spike regressors for flagged outlier volumes. To lower the computational load of whole-brain searchlight analyses, functional images were resampled to 3mm isotropic voxels and were restricted to a brain mask excluding the cerebellum. All ROI analyses were conducted using the original 2mm data resolution.

### Data Analysis

#### Behavioural Analysis

We first characterized the overall distribution of inter-event relational ratings in each group. To this end, the proportion of responses at each rating level (1–7) was computed per participant, yielding a 7-element profile vector per participant. Each group’s mean profile was then obtained by averaging these vectors across its participants. Qualitative evaluation of the group-average rating patterns revealed heavily skewed distributions, in which the majority of ratings in both groups (Spoiled: 64.9%; Naïve: 61.8%) were assigned the lowest possible value (“irrelevant”). This response profile did not differ between groups. Group-level response profiles were compared using the Euclidean distance between the two group-mean profiles as the test statistic.

Significance was assessed via a two-tailed permutation test in which participants’ group assignment were randomly shuffled and the group-mean distance was recomputed. This procedure was repeated 10,000 times creating a null distribution of distances. Comparing the unshuffled distance to this distribution yielded a p-value of 0.282, indicating that the overall response profile was shared across groups.

The severe skewness toward the lower end of the scale introduces challenges in statistical analysis (e.g. reduced sensitivity of measures of central tendency and dispersion, increased number of ties in correlation calculations). In the next sections, we report analyses designed to overcome these challenges, while also reporting the results of conventional approaches in order to provide full transparency and demonstrate the robustness of our findings. All main statistical inferences were implemented using non-parametric permutation tests. Permutation tests make no assumptions about the distribution of the data, and are particularly robust, as they generate empirical null distributions directly from the observed data. Because the same analysis is applied to both the observed and permuted datasets, any systematic bias of the chosen statistic is represented equally in the observed and null distributions^51, 52^. Finally, all inferential tests were based on participant-level summary statistics, preserving the participant as the unit of inference and avoiding inflation of degrees of freedom.

#### Group differences in rating distributions

We first aimed to test our preregistered hypothesis that differences in prior knowledge of the twist would alter the distributions of individual participants’ relational ratings, as reflected in their central tendency and dispersion. Although medians are often preferred as a descriptive statistic for ordinal data, the extreme concentration of responses at the lowest rating rendered the median insensitive to intersubject differences, as the medians tend to converge on the modal value. The mean, which is more sensitive to the shape of the distribution, was therefore a more suitable descriptive statistic for quantifying more subtle group differences in this case. For visualization purposes, group-averaged rating matrices were constructed by averaging each participant’s 666 scene-pair ratings within each condition, producing a lower triangular matrix per group in which each cell represented the mean relational rating assigned to each past–current scene pair. To quantify the overall similarity in the narrative structure captured by each group, Spearman’s correlation was computed between the two group-averaged matrices across all 666 scene-pair cells (participant-level analysis will be described in the section *intersubject similarity of ratings*).

To statistically test for group differences in means, the mean across all 666 ratings was calculated for each participant. These means were further averaged within the group, and the between-group difference in means was defined as a statistic in a permutation test. Under the null hypothesis of no group difference in means, participant labels are exchangeable across groups. Thus, across 10,000 iterations, the means of single participants were randomly assigned to 2 groups, and a random group average and between-group difference in averages were calculated. This procedure resulted in 10,000 random differences that constructed the null distribution, and the position of the true test statistic in relation to this distribution was used to determine the two-tailed P value of the test. For transparency, a similar test was used based on single-subject medians.

To quantify group differences in the dispersion of ratings, each participant’s variance was calculated across the 666 ratings. Variance was used as the measure of dispersion to maximize sensitivity to differences across the full rating range (interquartile range is additionally reported). Group differences in variances were assessed using a two-tailed permutation test, as described above.

#### Intersubject similarity of ratings

Following our second preregistered hypothesis, we tested whether ratings were more similar within groups than between groups. Given the extreme concentration of responses at the lowest rating level, conventional correlation measures can become dominated by ties and restricted variance. We therefore quantified inter-subject agreement in ratings using quadratic-weighted Cohen’s kappa coefficient. Quadratic-weighted Cohen’s kappa is particularly suitable for ordinal data because it is based on a confusion matrix, and disagreements are weighted according to their magnitude. As not all participants used all rating levels (resulting in sparse confusion matrices) and to control for idiosyncratic differences in scale use across participants, ratings were collapsed into three ordinal categories: low importance (1–2), moderate importance (3–5), and high importance (6–7). Intersubject agreement was then calculated as pairwise Cohen’s kappa for all participant pairs across both groups. For each participant, within-group similarity was defined as the mean agreement with all other members of the same condition, whereas between-group similarity was defined as the mean agreement with all members of the opposite condition. Group differences were assessed using one-tailed permutation tests (matching our directional preregistered hypothesis), with the test statistic defined as the difference between the mean within-group agreement (Spoiled+Naïve) and the mean between-group agreement. This main analysis was then broken down into its components by comparing within-Spoiled versus between-group agreement, within-Naïve versus between-group agreement, and within-Spoiled versus within-Naïve agreement. Each test was carried out by repeating the same analytic procedure after permuting participant labels, as previously described. For transparency, we additionally tested group differences in intersubject similarity using Pearson correlation on the full 1–7 rating scale.

### fMRI Analysis

#### fMRI Representations

Representation extraction followed the exact same analytical pipeline previously reported^37^. In short, a searchlight approach was employed, with spheres of 3-voxel (9 mm) radius confined by a standard MNI brain mask, excluding the cerebellum. In each sphere, representations of event boundaries were defined as the voxel pattern in the single TR that ends each scene. To separate the fMRI BOLD signals at event boundaries from signals of adjacent scenes, representations of scenes were defined as the average voxel pattern across all within-scene TRs that were at least 5 TRs (7.5 seconds) remote from adjacent event boundaries. The selection of a 5 TR gap was based on the estimation that the canonical HRF peaks at 4–6 seconds (3–4 TRs) of stimulus onset. Across all analyses, fMRI BOLD responses were shifted by 4 TRs (6 seconds) relative to the experimental stimuli to account for the canonical HRF delay.

#### Whole-brain Reactivation Analysis

To replicate our previous reactivation results, we again followed the exact same established analytic pipeline^37^, which will be summarized here. First, to detect similarities in neural representations indicative of reactivation, Pearson correlations were calculated between event boundary representations and scene representations for each searchlight sphere. Throughout all analyses, a leave-one-out between-participant (intersubject) approach was used. Specifically, event boundary representations were averaged across all participants of the same experimental group except one, and correlated with the scene representations extracted from that held-out participant. This was repeated for each participant in turn, and all correlations were Fisher z-transformed.

This procedure yielded event boundary × scene similarity matrices, in which the lower triangular part reflected correlations between each event boundary and its preceding scenes, and the upper triangular part reflected correlations between each event boundary and its following scenes (Fig. 1c, left). While these correlation matrices can provide a measure for reactivation of past scenes at event boundaries, they can also reflect inter-scene similarities. Yet, because inter-scene similarity contributes symmetrically to the upper and lower triangular parts of the similarity matrix, contrasting past and future correlations removes this symmetric component, allowing us to control perfectly for this potential confound. We therefore defined a reactivation index from this event boundary × scene similarity matrix, as the difference between the mean correlation of the lower triangular part (past scenes) and the mean of the upper triangular part (future scenes):

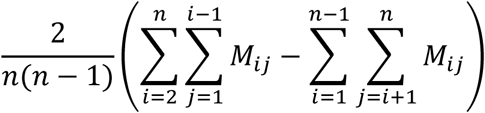

Where M is an n X n similarity matrix, i and j are row and column indices, respectively.

Thus, a positive reactivation index indicates greater reactivation of past compared to future events, and vice versa^37^.

For each participant and searchlight sphere, this reactivation index was computed and assigned to the anatomical coordinates of the sphere center, producing a single-subject whole-brain reactivation map. These single-subject maps of each experimental group were submitted to a voxel-wise one-sample one-tailed t-test, reflecting our directional hypothesis that reactivation would be greater for past than future scenes (i.e. positive reactivation indices, replicating our previous results^37^). Group maps were thresholded and corrected for multiple comparisons using cluster-mass permutation inference implemented in FSL’s randomise (cluster-forming threshold of *p < 0.001* and whole-brain family-wise error correction at *p < 0.05*). Between-group differences across the brain were assessed using the same procedure with a two-sample two-tailed t-test. Corrected maps were projected onto an MNI152 cortical surface for visualization using the Connectome Workbench.

### ROI Analysis

#### ROI Definition

Here we used three bilateral DMN Regions Of Interest (ROIs), previously shown to exhibit significant and replicable reactivation of past events at narrative event boundaries across two independent datasets^37^: hippocampus (HPC, 3,109 voxels), precuneus/retrosplenial cortex (PCUN, 7,947 voxels), and angular gyrus/lateral occipital cortex (ANG, 12,819 voxels). To assess effects across the DMN as a whole, voxels from these ROIs were merged into a composite DMN ROI. Throughout the paper, effects were primarily tested in this DMN ROI. Individual ROIs were analyzed subsequently to identify the sources of any overall DMN effect, with Holm-Bonferroni correction applied to correct for multiple comparisons.

#### Reactivation Analysis

For each participant and ROI, the representations of event boundaries and scenes were extracted, and reactivation indices were computed as described above. Group statistics were carried out using permutation tests, with the group-averaged reactivation index defined as the test statistic. Under the null hypothesis of no reactivation, the sign of single-subject reactivation indices could be flipped. Thus, for each group and ROI, reactivation indices of single participants were flipped randomly, and a random group mean was calculated. This procedure was repeated 10,000 times, resulting in 10,000 random means that constructed the null distribution. The position of the true test statistic in relation to the null distribution was used to determine the one-tailed p-value of the test. Between-group statistics were carried out using a two-tailed label-shuffle permutation test, as described above.

#### Reactivation Patterns

As described above, to isolate the asymmetric component of neural reactivation, the reactivation index contrasted the mean correlation between representations of event boundaries and preceding scenes (lower triangular part of the similarity matrix) with the corresponding mean correlation between event boundaries and subsequent scenes (upper triangular part of the similarity matrix). This procedure collapses the full event boundary × scene similarity matrix into a single value per brain region. However, because the difference between two means is mathematically equivalent to the mean of their pairwise differences, the reactivation index can be decomposed into boundary-scene-specific contributions while preserving the granular structure of the matrix. Specifically, for each event boundary i and preceding scene j (i>j), we defined a granular reactivation index as the difference between each matrix entry and its symmetric counterpart (Fig. 1c, right). The reactivation pattern was then defined as the set of all such values:

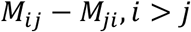

Where M is the EB X scene similarity matrix and i and j are row and column indices, respectively.

This reactivation pattern consisting of 666 granular reactivation values for each subject and ROI, reflecting the extent to which each scene was reactivated at each event boundary, and thereby providing a neural measure directly comparable to the behavioural relational ratings (Fig. 1b, c).

#### Dispersion of the reactivation patterns

The variance across the granular reactivation pattern was computed for each participant and ROI. Between-group statistics were obtained using a two-tailed label-shuffle permutation test, as described above.

#### Intersubject similarity of reactivation patterns

Within-group similarity of the reactivation pattern was computed using a leave-one-out approach: each participant’s pattern was Spearman correlated with the mean pattern of all other members of their own group. Between-group similarity was computed as each participant’s correlation with the mean pattern of all members of the opposite group. Statistical significance was assessed using a label-shuffle permutation test, in which the test statistic was the difference between mean similarity values: Our primary analysis was within-group (Spoiled+Naïve) versus between-group similarity. Subsequent breakdown of this analysis included within-Spoiled versus between-group similarity, within-Naïve versus between-group similarity, and within-Spoiled versus within-Naïve similarity. On each of 1,000 iterations, participants were randomly assigned to groups of the same original sizes, event boundary × scene similarity matrices were computed, reactivation patterns were derived, and intersubject similarities were computed within and between groups, yielding a null distribution for each comparison. The tests’ true statistics were compared to these distributions to derive p values (one-tailed for the within vs. between comparison, reflecting our directional hypothesis of greater within- than between-group similarity; two-tailed for the within group similarity comparison).

#### Brain–Behaviour correspondence

We next tested whether neural reactivation in DMN regions was positively associated with the relational ratings of scene-pairs, as measured in our behavioural experiment (i.e. higher reactivation of scenes that are judged as more important for understanding each current event, and vice versa). However, the heavily skewed behavioural ratings produced extensive tied ranks, meaning that the majority of scene pairs were indistinguishable in rank and therefore contributed limited information to a direct correlation across the 666 values. For this reason, we estimated the mean reactivation associated with each rating level and assessed the monotonic relationship across the seven rating categories. Specifically, for each behavioural-fMRI participant pair of the same experimental condition, the 666 granular reactivation values were grouped according to the corresponding behavioural rating (1–7). Mean reactivation was then computed separately for each rating level, yielding seven mean reactivation values. Spearman’s correlation was computed between the seven rating levels and their corresponding mean reactivation values. This binning approach ensured that each rating level contributes a single noise-reduced estimate of mean reactivation, reducing the influence of unequal frequencies across rating levels and extensive tied ranks. For each behavioural participant, the correlation coefficients with all fMRI participants of the same experimental condition were then averaged, yielding a single value per behavioural participant per ROI. Group statistics were obtained using a sign-flip permutation test, under the null hypothesis of no relation between behavioural and neural measures, and between-group differences were assessed using a label-shuffle permutation test, as described above. To determine whether the results were robust to the direction of averaging, this analysis was repeated by averaging the correlation coefficients across behavioural participants for each fMRI participant within the same condition.

As a robustness check, we repeated the analysis using Spearman’s correlation on the raw values rather than the binned data. For each behavioural-fMRI participant pair from the same condition, Spearman’s correlation was computed between the 666 behavioural ratings and the corresponding 666 granular reactivation values, and the same statistical procedure described above was applied to these correlation values.

To test whether the hippocampal group difference in brain–behaviour coupling could be explained by differences in the properties of reactivation or rating distributions, we fit two ordinary least squares regression models predicting hippocampal brain–behaviour coupling from condition together with covariates capturing the mean and variance of the neural or behavioural measures. Model 1 was defined at the neural-subject level (*n = 57*): coupling ∼ condition + mean neural reactivation + neural reactivation variance, using the correlation coefficients computed for each fMRI participant, as described above. Model 2 was defined at the behavioural-subject level (*n = 95*): coupling ∼ condition + mean behavioural rating + behavioural rating variance, using the correlation coefficients computed for each behavioural participant. For each model, the statistical significance of the condition coefficient was assessed using a two-tailed label-shuffle permutation test: condition labels were randomly reassigned across participants (10,000 iterations), the regression was refit for the shuffled data of each permutation, and the position of the true condition coefficient relative to this null distribution determined the two-tailed P-value.

Because the behavioural rating patterns were highly similar across groups (*r = 0.935* between group-average ratings), whole-matrix behavioural–neural correlations were expected to be similar across within-group and between-group pairings. To confirm this, behavioural ratings were correlated with neural reactivation patterns from participants in the other experimental group using the same statistical procedures described above.

To assess whether neural reactivation reflected behavioural differences between groups, we focused on the subset of scene pairs that showed the largest behavioural divergence between conditions. For each of the 666 scene pairs, a Welch’s t-test compared Spoiled and Naïve behavioural ratings. The top 25% of scene pairs (166 pairs) showing the largest absolute between-group differences were selected. We then tested whether the corresponding granular reactivation values also differed between groups. For each fMRI participant and ROI, the corresponding granular reactivation values were averaged across these selected scene pairs, yielding a single mean reactivation value. These values were then analyzed using within- and between-group permutation tests, as described above, to determine whether reactivation was significant within each group, and whether its magnitude differed between groups. Note that because scene pairs were selected exclusively on the basis of the behavioural data, the neural data remained statistically independent of the selection procedure. To assess the robustness of the findings to the choice of selection threshold, the analysis was repeated using the top 20% and 30% of scene pairs showing absolute between-group behavioural difference.

To determine whether this effect could be explained by the Naïve group’s overall higher reactivation, for each participant and ROI, we computed the difference between mean reactivation across the selected Top-N scene pairs and mean reactivation across the remaining non-selected pairs. This within-participant contrast removes additive differences in overall reactivation level. We then tested for a group × selection interaction by comparing these differences between Naïve and Spoiled participants using a one-tailed label-shuffle permutation test, as described above. This analysis therefore tested whether the between-group difference in reactivation was disproportionately expressed for behaviourally divergent scene pairs rather than reflecting a general elevation of reactivation in the Naïve group. The analysis was conducted for the top 25% of scene pairs, with the top 20% and 30% examined as robustness analyses.

For visualization purposes only, behavioural ratings were averaged across the selected pairs within each participant, and the resulting group differences were plotted. Scene pairs were selected based on the absolute magnitude of the behavioural group difference, irrespective of direction. The signed behavioural difference within this selected subset was therefore examined descriptively to characterize the direction of the selected effects. Because the same behavioural data were used both to select the scene pairs and to quantify the behavioural group difference, this comparison was not treated as an independent inferential test.

## Acknowledgments

This study was supported by the Royal Society’s Dorothy Hodgkin Fellowship DHF\R1\241022 (A.H.); and the European Union under a European Research Council Starting Grant awarded to A.B.-Y. (101077119-MEME). Views and opinions expressed are however those of the authors only and do not necessarily reflect those of the European Union or the European Research Council Executive Agency. Neither the European Union nor the granting authority can be held responsible for them.

---

**Supplementary Figure 1.**
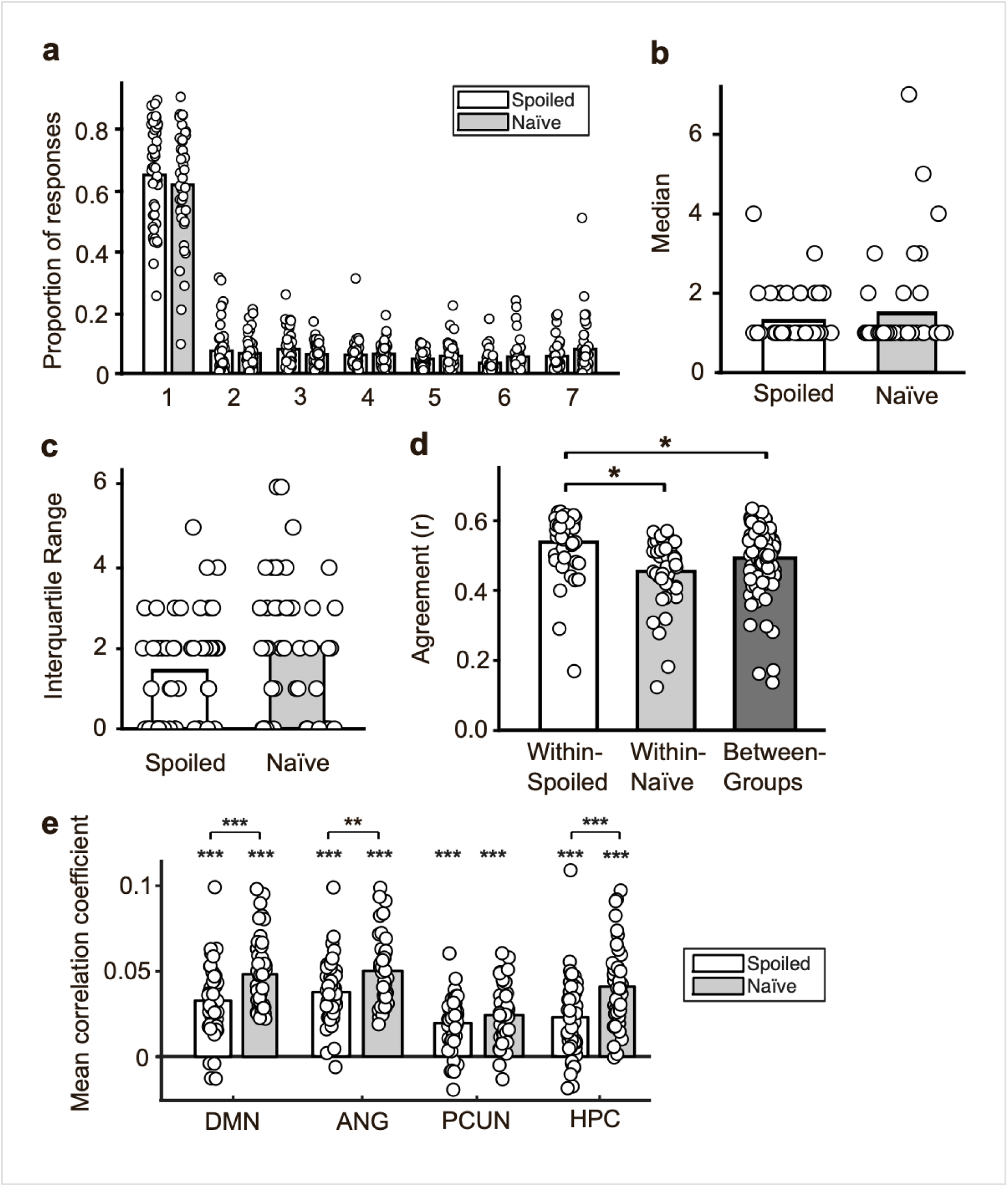
Rating distributions and robustness analyses. (a) Proportion of responses assigned to each rating level (1–7) in each group (Spoiled: white bars, n = 50; Naïve: gray bars, n = 45). The distributions were heavily right-skewed in both groups, with approximately 65% of responses assigned a rating of 1. (b) Median of ratings for each participant across all 666 scene pairs. The Naïve group showed a non-significant higher median compared to the Spoiled group (Spoiled: *M = 1.30*, *SD = 0.61*; Naïve: *M = 1.49*, *SD = 1.22*; *p = 0.408*, two-tailed permutation test). (c) Interquartile range of ratings for each participant across all 666 scene pairs. The Naïve group showed a non-significant higher IQR compared to the Spoiled group (Spoiled: *M = 1.44*, *SD = 1.33*; Naïve: *M = 2.02*, *SD = 1.73*; *p = 0.073*, two-tailed permutation test). (d) Pearson intersubject agreement of ratings within and between groups, averaged across all pairs for each participant. Within-Spoiled agreement (*M = 0.520*, *SD = 0.082*) significantly exceeded both between-group agreement (*M = 0.475*, *SD = 0.090*; *p = 0.0117*, one-tailed permutation test) and within-Naïve agreement (*M = 0.439*, *SD = 0.092*; *p = 0.0272*, two-tailed permutation test), while within-Naïve agreement did not differ from between-group (*p = 0.9805*, one-tailed). (e) Spearman correlations were computed between raw behavioural ratings and corresponding granular neural reactivation values, for each behavioural–fMRI participant pair from the same experimental condition (Spoiled: white bars; Naïve: gray bars). Both groups showed significant positive correlations across all ROIs (all p < 0.001, one-tailed permutation test). Between-group comparisons showed significantly stronger coupling in the Naïve group in DMN (*p < 0.001*), angular gyrus (*p = 0.0048*), and hippocampus (*p < 0.001*), but not precuneus (*p = 0.4968*, two-tailed permutation tests). Note that correlation coefficients were substantially smaller than the correlation shown in Fig. 5a, consistent with the expected sensitivity advantage of the binning approach when ratings are heavily skewed and contain extensive tied ranks. Bars indicate group means; dots represent individual participants. Asterisks above the bars indicate that the corresponding group means differ significantly from zero (sign-flip permutation test, one-tailed); bracket asterisks indicate significant between-group differences (label-shuffle permutation test, two-tailed). *\*p < 0.05, **p < 0.01, ***p < 0.001*.

**Supplementary Figure 2.**
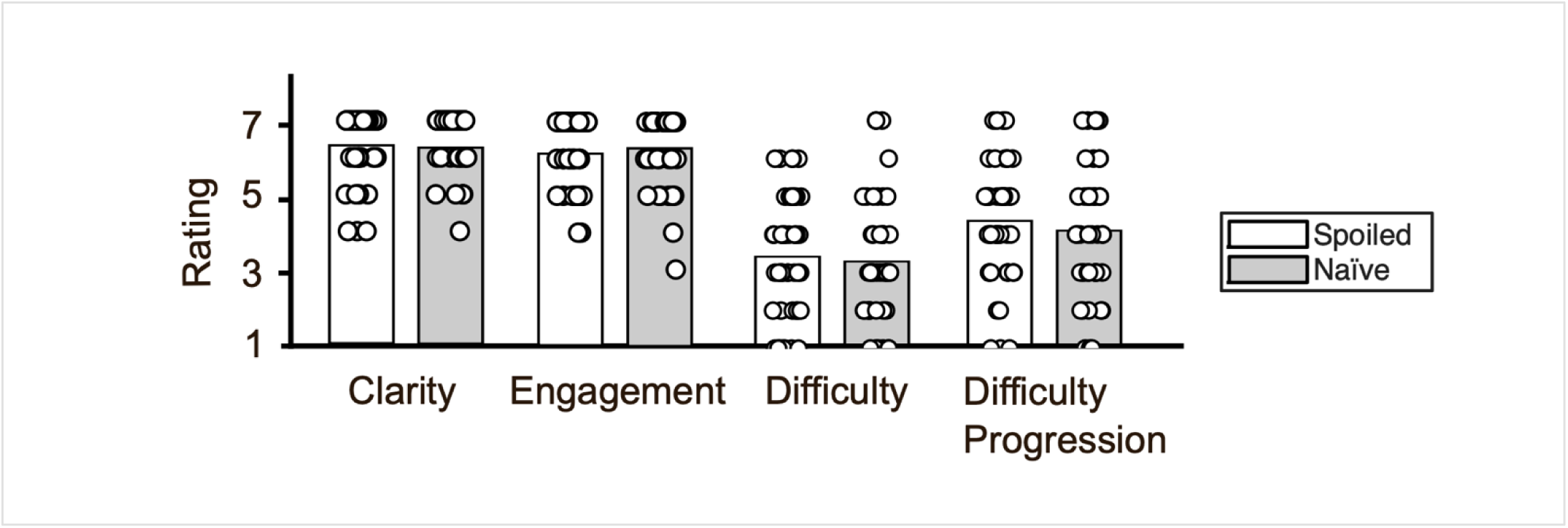
Post-experiment task assessments. Post-experiment ratings for four dimensions of task experience: clarity of instructions, engagement with the film, perceived difficulty, and perceived progression of difficulty over the course of the film, shown separately for the Spoiled (white bars; *n = 50*) and Naïve (gray bars; *n = 45*) groups. Ratings were made on a 1–7 scale. Both groups rated the instructions as clear (Spoiled: *M = 6.44*, *SD = 0.88*; Naïve: *M = 6.38*, *SD = 1.11*) and the movie as engaging (Spoiled: *M = 6.26*, *SD = 0.88*; Naïve: *M = 6.40*, *SD = 0.96*); task difficulty was moderate (Spoiled: *M = 3.42*, *SD = 1.55*; Naïve: *M = 3.29*, *SD = 1.63*), with a slight perceived increase over time (Spoiled: *M = 4.40*, *SD = 1.50*; Naïve: *M = 4.13*, *SD = 1.75*). No significant between-group differences were observed for any dimension (two-tailed permutation tests; clarity: *p = 0.850*; engagement: *p = 0.502*; difficulty: *p = 0.696*; difficulty progression: *p = 0.451*). Bars indicate group means; dots represent individual participants.

**Supplementary Figure 3.**
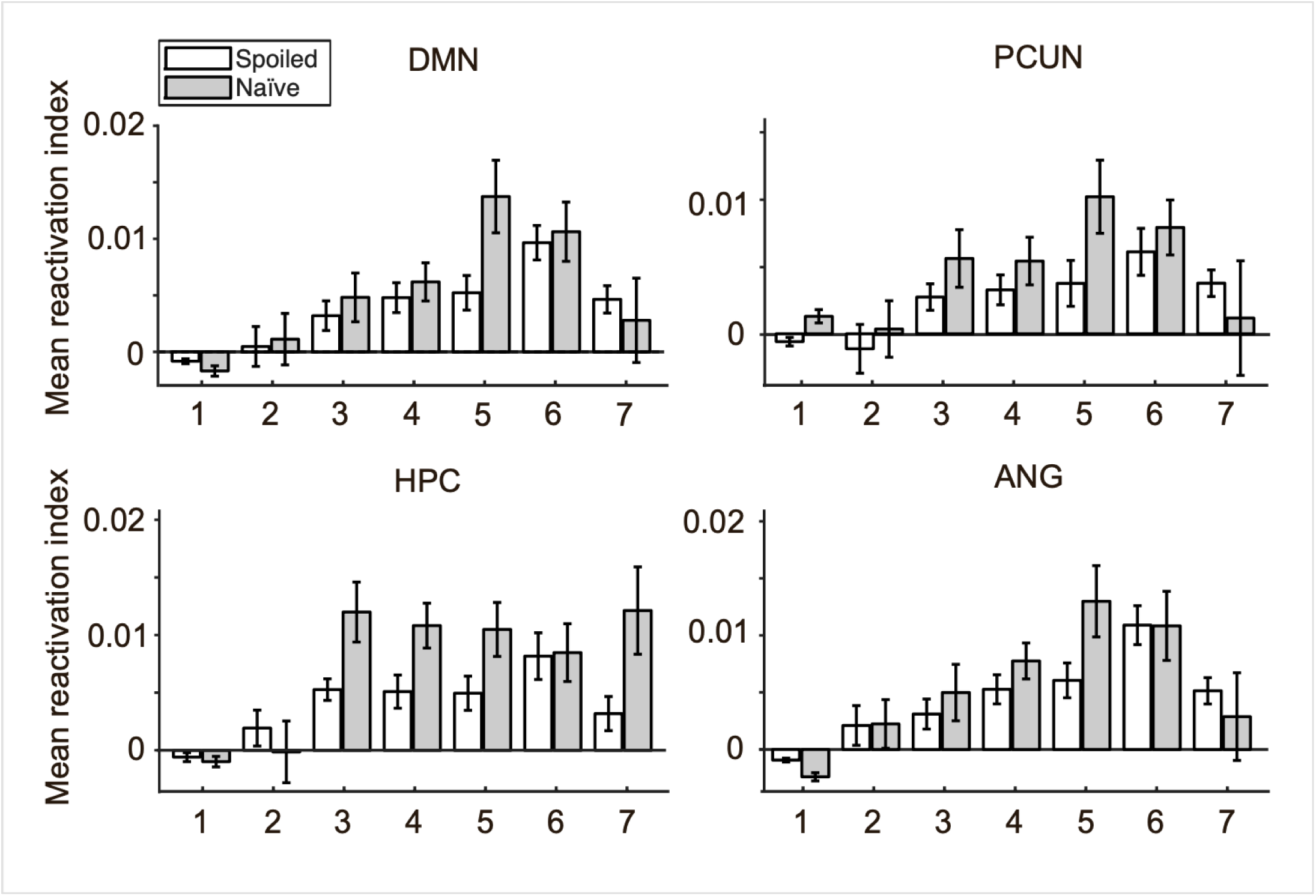
Neural reactivation as a function of relational rating. Mean reactivation index per rating level (1–7) shown separately for the Spoiled (white bars; *behavioural n = 50*, *fMRI n = 18*) and Naïve (gray bars; *behavioural n = 45*, *fMRI n = 39*) groups across ROIs. For each group and rating level, mean reactivation was first computed for every behavioural–fMRI participant pair and then averaged across all same-condition participant pairs. This descriptive figure illustrates the binned data underlying the monotonic brain–behaviour relationship summarized by the Spearman correlations in Fig. 5a. Error bars indicate SEM.

**Supplementary Table 1.** Cluster statistics for the within-group whole-brain reactivation of past scenes at event boundaries. Clusters surviving whole-brain correction (cluster-forming threshold *p < 0.001*; cluster-mass FWE-corrected *p < 0.05*, permutation tests). MNI coordinates correspond to the peak voxel of each cluster. Anatomical labels are based on the Harvard-Oxford Cortical/Subcortical Structural Atlases (FSL).

| Group | #Voxels | Peak t | p (FWE-corr) | MNI x | MNI y | MNI z | Region |
|---|---|---|---|---|---|---|---|
| Spoiled | 1954 | 12.95 | <0.001 | 30 | -75 | -6 | R Occipital Fusiform Gyrus; cluster extends into ANG/LOC, PCUN, and HPC |
| Spoiled | 1044 | 12.54 | <0.001 | -30 | -54 | -3 | L Lingual Gyrus; cluster extends into ANG/LOC, PCUN, and HPC |
| Spoiled | 45 | 5.51 | 0.049 | -12 | -96 | 18 | L Occipital Pole |
| Naïve | 8815 | 21.99 | <0.001 | -21 | -60 | -3 | L Lingual Gyrus; cluster extends bilaterally through ANG/LOC, PCUN, HPC |
| Naïve | 259 | 7.04 | 0.005 | 33 | 15 | 36 | R Middle Frontal Gyrus |
| Naïve | 98 | 5.16 | 0.013 | 48 | 42 | 12 | R Frontal Pole |
| Naïve | 64 | 5.95 | 0.021 | 15 | 54 | 21 | R Frontal Pole |
| Naïve | 62 | 5.15 | 0.026 | 9 | 9 | 60 | R Superior Frontal Gyrus |
| Naïve | 55 | 4.81 | 0.037 | -48 | 27 | 36 | L Middle Frontal Gyrus |
| Naïve | 53 | 5.1 | 0.037 | 0 | 18 | 24 | Anterior Cingulate Gyrus |
| Naïve | 50 | 6.86 | 0.034 | 0 | 45 | -6 | Paracingulate Gyrus |

**Supplementary Table 2.** Cluster statistics for the between-group differences in whole-brain reactivation. Clusters surviving whole-brain FWE correction (cluster-forming threshold *p < 0.001*; cluster-mass FWE-corrected *p < 0.05*, permutation tests). MNI coordinates correspond to the peak voxel of each cluster. Anatomical labels are based on the Harvard-Oxford Cortical/Subcortical Structural Atlases (FSL).

| Contrast | #Voxels | Peak t | p (FWE-corr) | MNI x | MNI y | MNI z | Region |
|---|---|---|---|---|---|---|---|
| Spoiled > Naïve | 155 | 7.13 | <0.001 | 69 | -27 | 9 | R Superior Temporal Gyrus, posterior division |
| Spoiled > Naïve | 45 | 6.58 | 0.033 | -60 | -18 | 3 | L Planum Temporale |
| Spoiled > Naïve | 43 | 5.01 | 0.046 | 51 | 15 | -24 | R Temporal Pole |
| Naïve > Spoiled | 301 | 5.8 | <0.001 | -12 | -48 | -6 | L Lingual Gyrus; cluster extends into HPC and PCUN |
| Naïve > Spoiled | 182 | 5.93 | 0.002 | 6 | -69 | 45 | Precuneus |
| Naïve > Spoiled | 55 | 4.82 | 0.016 | -21 | -48 | 60 | L Superior Parietal Lobule |
| Naïve > Spoiled | 51 | 5.54 | 0.026 | -24 | 6 | 60 | L Superior Frontal Gyrus |

**Supplementary Table 3.**
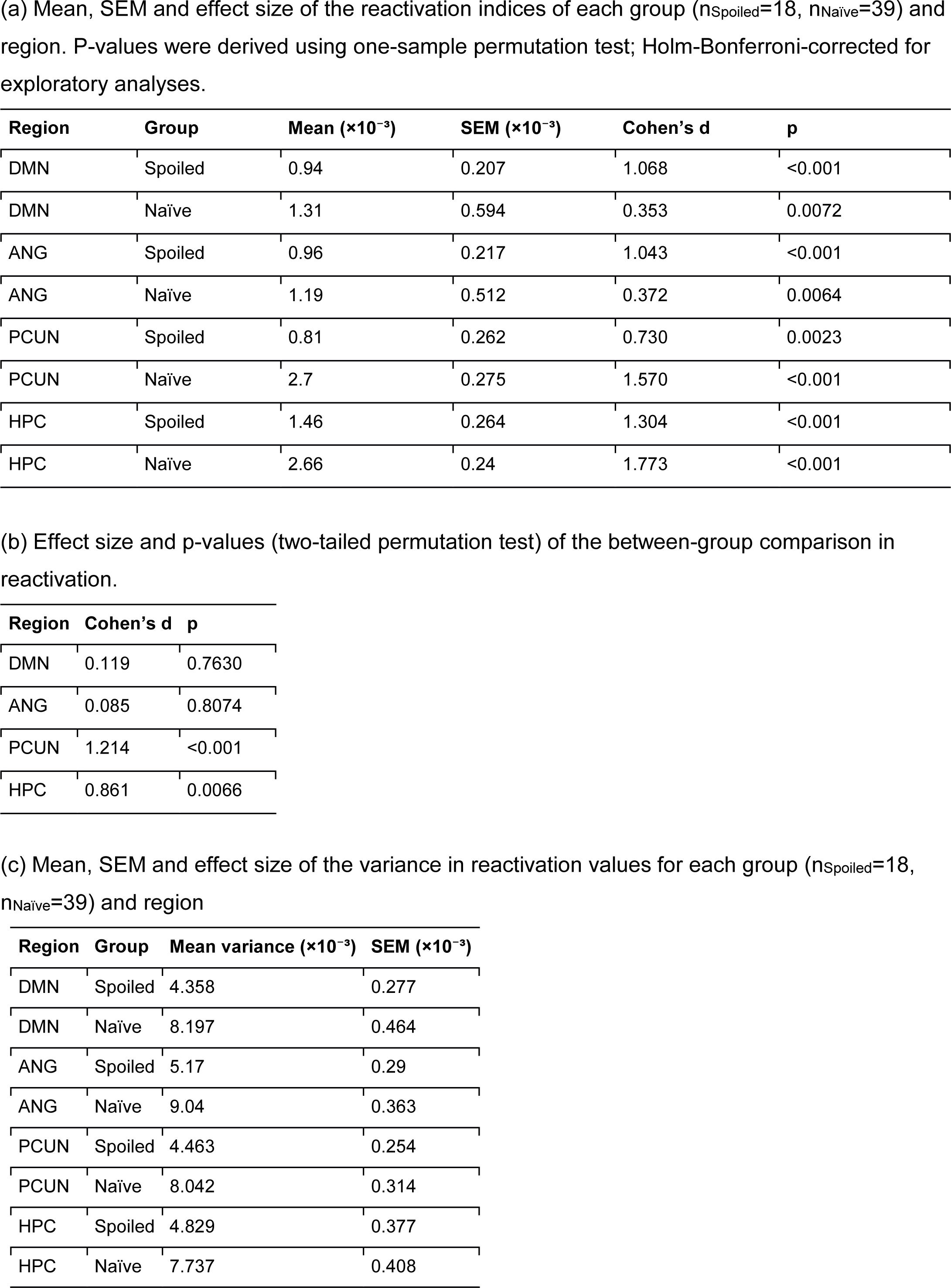
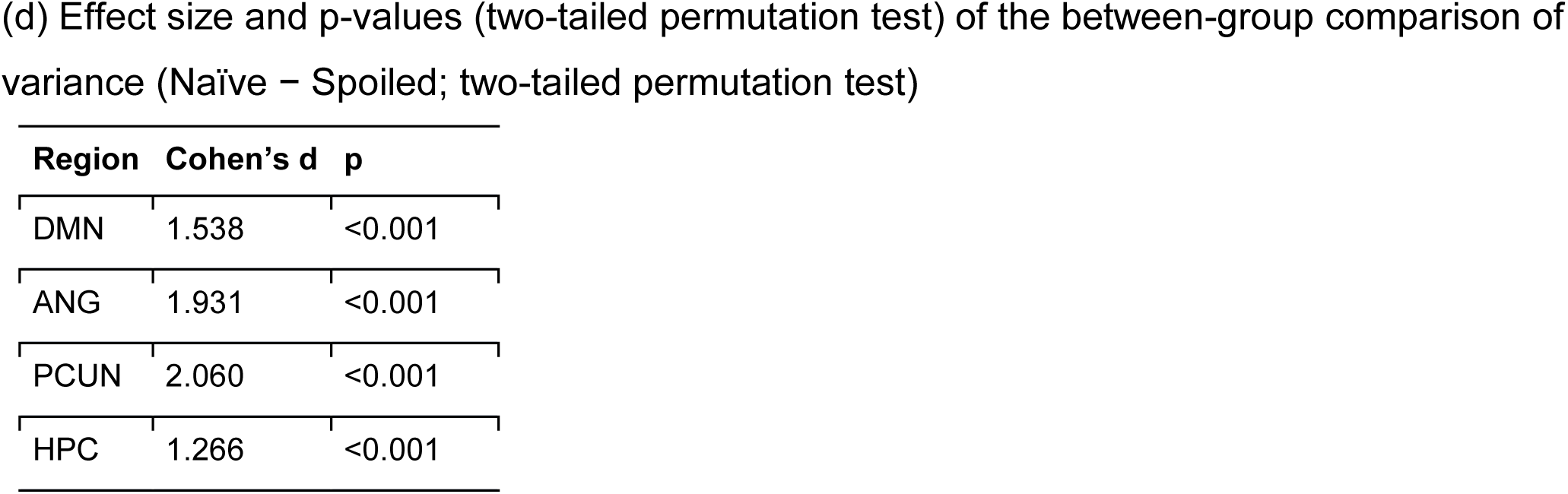
ROI reactivation index and variance statistics.

| **Region** | **Group** | **Mean (×10<sup>−3</sup>)** | **SEM (×10<sup>−3</sup>)** | **Cohen’s d** | **p** |
|---|---|---|---|---|---|
| DMN | Spoiled | 0.94 | 0.207 | 1.068 | <0.001 |
| DMN | Naïve | 1.31 | 0.594 | 0.353 | 0.0072 |
| ANG | Spoiled | 0.96 | 0.217 | 1.043 | <0.001 |
| ANG | Naïve | 1.19 | 0.512 | 0.372 | 0.0064 |
| PCUN | Spoiled | 0.81 | 0.262 | 0.730 | 0.0023 |
| PCUN | Naïve | 2.7 | 0.275 | 1.570 | <0.001 |
| HPC | Spoiled | 1.46 | 0.264 | 1.304 | <0.001 |
| HPC | Naïve | 2.66 | 0.24 | 1.773 | <0.001 |
| **Region** | **Cohen’s d** | **p** |  |  |  |
| DMN | 0.119 | 0.7630 |  |  |  |
| ANG | 0.085 | 0.8074 |  |  |  |
| PCUN | 1.214 | <0.001 |  |  |  |
| HPC | 0.861 | 0.0066 |  |  |  |
| **Region** | **Group** | **Mean variance (×10<sup>−3</sup>)** | **SEM (×10<sup>−3</sup>)** |  |  |
| DMN | Spoiled | 4.358 | 0.277 |  |  |
| DMN | Naïve | 8.197 | 0.464 |  |  |
| ANG | Spoiled | 5.17 | 0.29 |  |  |
| ANG | Naïve | 9.04 | 0.363 |  |  |
| PCUN | Spoiled | 4.463 | 0.254 |  |  |
| PCUN | Naïve | 8.042 | 0.314 |  |  |
| HPC | Spoiled | 4.829 | 0.377 |  |  |
| HPC | Naïve | 7.737 | 0.408 |  |  |
| **Region** | **Cohen’s d** | **p** |  |  |  |
| DMN | 1.538 | <0.001 |  |  |  |
| ANG | 1.931 | <0.001 |  |  |  |
| PCUN | 2.060 | <0.001 |  |  |  |
| HPC | 1.266 | <0.001 |  |  |  |

**Supplementary Table 4.**
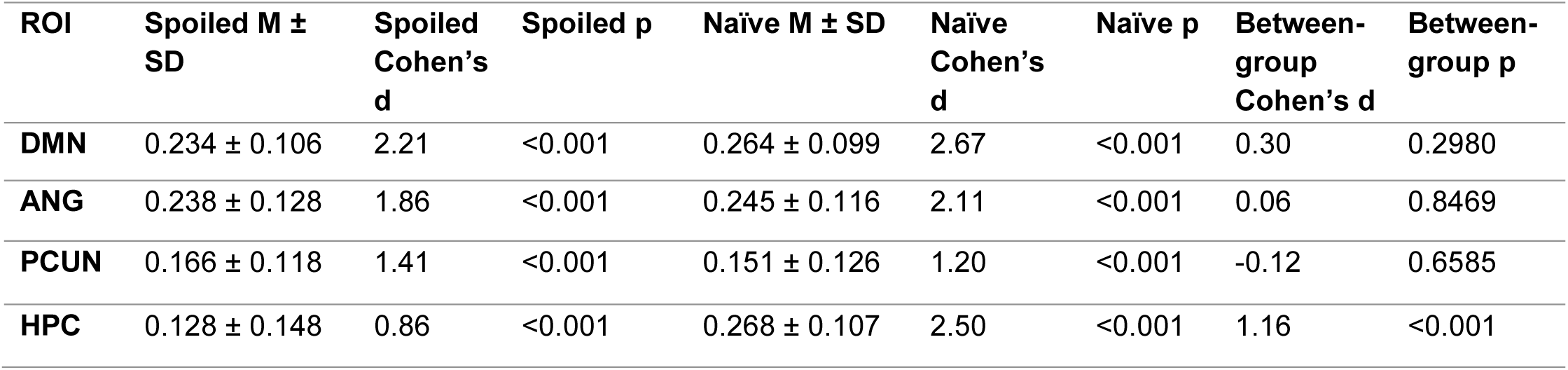
Brain–behaviour correspondence using fMRI-participant-level averaging. As a robustness check of the primary brain–behaviour analysis (see Methods; Fig. 5a), correlation coefficients were averaged across all behavioural participants for each fMRI participant within the same experimental condition, reversing the averaging procedure used in the primary analysis (i.e. averaging across fMRI participants per behavioural participant). ANG/PCUN/HPC p-values are Holm-Bonferroni-corrected. Within-group tests are one-tailed and between-group tests are two-tailed (permutation tests). This reverse-averaging approach reproduces the same pattern as the primary analysis, demonstrating that the observed brain–behaviour relationships were robust to the direction of averaging.

**Supplementary Table 5.** Brain-behaviour correlations within and across experimental conditions. For each ROI, Spearman correlations were computed between each behavioural participant’s ratings and neural reactivation patterns from fMRI participants in the same condition (Same) or the opposite condition (Cross). Within-condition correlations were assessed using one-tailed permutation tests; Same versus Cross differences were assessed using two-tailed permutation tests; p-values for ANG, PCUN, and HPC were Holm-Bonferroni corrected. Brain–behaviour coupling was significant in both Same and Cross pairings in every ROI, with no significant Same-vs-Cross difference, consistent with the high similarity between the behavioural rating patterns of the two groups (Fig. 2a).

| **ROI** | **Spoiled-Same r (p)** | **Cohen’s d** | **Spoiled-Cross r (p)** | **Cohen’s d** | **Δr (Same − Cross) (p)** | **Naïve-Same r (p)** | **d** | **Naïve-Cross r (p)** | **Cohen’s d** | **Δr (Same − Cross) (p)** |
|---|---|---|---|---|---|---|---|---|---|---|
| DMN | 0.033 (<0.001) | 1.65 | 0.043 (<0.001) | 2.07 | 0.061 | 0.048 (<0.001) | 2.33 | 0.039 (<0.001) | 2.20 | 0.104 |
| ANG | 0.038 (<0.001) | 2.07 | 0.045 (<0.001) | 2.33 | 0.4138 | 0.050 (<0.001) | 2.55 | 0.043 (<0.001) | 2.58 | 0.5126 |
| PCUN | 0.019 (<0.001) | 1.25 | 0.022 (<0.001) | 1.46 | 0.5711 | 0.024 (<0.001) | 1.45 | 0.022 (<0.001) | 1.47 | 0.7471 |
| HPC | 0.023 (<0.001) | 1.03 | 0.035 (<0.001) | 1.53 | 0.111 | 0.041 (<0.001) | 1.75 | 0.028 (<0.001) | 1.37 | 0.105 |

**Supplementary Table 6.** Robustness of the brain-behaviour correspondence of divergent-scene-pairs across selection thresholds. Behavioural ratings and neural reactivation were averaged within each participant over the N scene pairs showing the largest absolute between-group difference in behavioural ratings (Welch t-test, ranked by |t|). Two selection thresholds, 20% and 30% of the 666 scene pairs are shown (the Top 25% threshold, is reported in the main text; Fig. 5b). Behavioural means are presented descriptively, because the same behavioural data were used to select the scene pairs. In (a), neural between-group comparisons were performed using the same two-tailed permutation tests as in the main analysis, ANG, PCUN, and HPC, p-values are Holm-Bonferroni-corrected. Neural reactivation was consistently stronger in the Naïve group than in the Spoiled group across all thresholds and all regions, confirming that scene pairs distinguished by the largest behavioural rating differences also show the largest neural differences between groups; statistical significance for individual ROIs varied somewhat with threshold size but the direction of the effect remains. (b) and (c) report a companion analysis (see Methods) testing whether this reactivation difference was specific to the selected pairs, rather than driven by a generic between-group difference in reactivation.

| Threshold | Behaviour (Spoiled / Naïve) | DMN (Spoiled / Naïve) | DMN Cohen's d (p) | ANG (Spoiled / Naïve) | ANG Cohen's d (p) | PCUN (Spoiled / Naïve) | PCUN Cohen's d (p) | HPC (Spoiled / Naïve) | HPC Cohen's d (p) |
|---|---|---|---|---|---|---|---|---|---|
| **Top 20% (133 pairs)** | 1.86 ± 0.59 / 2.50 ± 1.13 | 7.07 ± 2.76 / 10.19 ± 8.15 | 0.45 (0.123) | 6.01 ± 3.01 / 10.40 ± 7.30 | 0.70 (0.033) | 6.13 ± 2.85 / 9.68 ± 7.87 | 0.53 (0.069) | 9.59 ± 2.79 / 15.28 ± 7.66 | 0.87 (0.011) |
| **Top 30% (200 pairs)** | 1.90 ± 0.55 / 2.43 ± 1.04 | 6.73 ± 2.71 / 10.51 ± 6.75 | 0.65 (0.024) | 6.41 ± 2.74 / 11.27 ± 6.57 | 0.86 (0.010) | 5.67 ± 2.75 / 9.97 ± 5.74 | 0.86 (0.010) | 8.81 ± 3.13 / 12.31 ± 5.60 | 0.70 (0.017) |
| (b) Mean and SEM of the reactivation difference between the selected and non-selected scene pairs for each group and region, and within-group p-values and effect sizes testing whether this difference was significantly above zero (one-tailed sign-flip permutation test; ANG, PCUN, and HPC Holm-Bonferroni-corrected, DMN uncorrected). |  |  |  |  |  |  |  |  |  |
| **Threshold** | **DMN (Spoiled / Naïve)** | **DMN Cohen's d (p)** | **ANG (Spoiled / Naïve)** | **ANG Cohen's d (p)** | **PCUN (Spoiled / Naïve)** | **PCUN Cohen's d (p)** | **HPC (Spoiled / Naïve)** | **HPC Cohen's d (p)** |  |
| **Top 20% (133 pairs)** | 7.66 ± 0.79 / 10.61 ± 1.12 | 2.29 (<0.001) / 1.52 (<0.001) | 6.31 ± 0.84 / 11.03 ± 1.04 | 1.77 (<0.001) / 1.70 (<0.001) | 6.64 ± 0.87 / 8.32 ± 1.32 | 1.79 (<0.001) / 1.01 (<0.001) | 10.16 ± 0.85 / 15.17 ± 1.26 | 2.83 (<0.001) / 1.93 (<0.001) |  |
| **Top 30% (200 pairs)** | 8.28 ± 0.89 / 12.51 ± 0.95 | 2.20 (<0.001) / 2.11 (<0.001) | 7.79 ± 0.97 / 13.71 ± 0.91 | 1.89 (<0.001) / 2.41 (<0.001) | 6.96 ± 0.92 / 9.85 ± 1.06 | 1.78 (<0.001) / 1.49 (<0.001) | 10.52 ± 1.03 / 13.15 ± 0.95 | 2.42 (<0.001) / 2.22 (<0.001) |  |
| (c) Between-group p-values (one-tailed permutation test) and effect sizes testing whether the reactivation difference in (b) was significantly larger in the Naïve than the Spoiled group; ANG, PCUN, and HPC Holm-Bonferroni-corrected, DMN uncorrected. |  |  |  |  |  |  |  |  |  |
| **Threshold** | **DMN Δ(Naïve−Spoiled)** | **DMN Cohen's d (p)** | **ANG Δ(Naïve−Spoiled)** | **ANG Cohen's d (p)** | **PCUN Δ(Naïve−Spoiled)** | **PCUN Cohen's d (p)** | **HPC Δ(Naïve−Spoiled)** | **HPC Cohen's d (p)** |  |
| **Top 20% (133 pairs)** | 2.96 | 0.48 (0.044) | 4.72 | 0.82 (0.007) | 1.68 | 0.23 (0.202) | 5.01 | 0.73 (0.012) |  |
| **Top 30% (200 pairs)** | 4.23 | 0.79 (0.003) | 5.92 | 1.13 (<0.001) | 2.89 | 0.49 (0.090) | 2.63 | 0.48 (0.090) |  |

